# Intracellular Energy Controls Dynamics of Stress-induced Ribonucleoprotein Granules

**DOI:** 10.1101/2022.03.28.486002

**Authors:** Tao Wang, Xibin Tian, Yura Jang, Paul Huang, Chan Hyun Na, Jiou Wang

## Abstract

Energy metabolism and membraneless organelles have been implicated in human diseases including neurodegeneration. How energy stress regulates ribonucleoprotein particles such as stress granules (SGs) is still unclear. Here we identified a unique type of granules formed under energy stress and uncovered the mechanisms by which the dynamics of diverse stress-induced granules are regulated. Severe energy stress induced the rapid formation of energy-associated stress granules (eSGs), whereas moderate energy stress delayed the clearance of conventional SGs. The formation of eSGs or the clearance of conventional SGs was regulated by the mTOR-4EBP1-eIF4E pathway or eIF4A1, involving eIF4F complex assembly or RNA condensation, respectively. In ALS patients’ neurons or cortical organoids, the eSG formation was enhanced, and conventional SG clearance was impaired. These results reveal a critical role for intracellular energy in the regulation of diverse granules and suggest that an imbalance in these dynamics may contribute to the pathogenesis of relevant diseases.

## Introduction

The assembly of ribonucleoprotein (RNP) granules, membraneless organelles composed of RNA and RNA-binding proteins (RBPs), via liquid-liquid phase separation (LLPS) is emerging as a principal mechanism for biochemical organization^1^. These structures play critical roles in multiple biological processes, including mRNA processing and translational control. The stress granule (SG) is a type of cytoplasmic RNP granule that is dynamically assembled in response to various stresses, such as oxidative, osmotic, or proteotoxic stress^2, 3^. SGs are composed of polyadenylated mRNA, stalled preinitiation complexes containing 40S ribosomal subunits and translation initiation factors, and various RBPs, including G3BP1/G3BP2, which have been proposed as the central scaffold proteins essential for multivalent interactions and SG assembly^2–4^.

Whereas transient SG formation in response to acute stress may be protective, disturbed SG dynamics such as its enhanced SG formation or reduced clearance have been thought to contribute to the pathogenesis of multiple neurodegenerative diseases (NDDs), including amyotrophic lateral sclerosis (ALS) and frontotemporal dementia (FTD), two related conditions with overlapping causes and pathologies^5–7^. For example, a predominant pathological hallmark of ALS/FTD is the intracellular aggregation of TDP-43, an RBP that serves as a building block of SGs^8^. An increasing number of the ALS/FTD-associated RBPs, including FUS, hnRNPA1, hnRNPA2B1, TIA1, Matrin 3, and Ataxin 2, are found as SG components^7, 9–12^; moreover, several ALS/FTD-associated non-RBPs, including dipeptide repeats (C9-DPRs) derived from *C9orf72* hexanucleotide repeat expansions, SOD1, p62, UBQLN2, and VCP, are found to co-localize with SGs and regulate their dynamics^13–17^. Forced SG formation by an optogenetic approach causes cytoplasmic inclusions that recapitulate the pathology of ALS-FTD^18^. Finally, inhibiting SG accumulation by targeting eIF2α phosphorylation, one of the most studied drivers for SG formation, or key SG proteins such as TDP-43, Ataxin 2, or TIA1 protects against disease-associated phenotypes in neuron culture or animal models of ALS/FTD^12, 19–22^.

To date, most studies of SGs have been based on their induction by exogenous stressors such as arsenite, but SGs induced by endogenous stressors remain largely unexplored. It has been suggested that cellular energy metabolism is closely related to SG dynamics. For example, mitochondrial poisoning or inhibition of glycolysis induces rapid SG formation in mammalian cells ^23–25^. Moreover, chronic glucose starvation in combination with depletion of amino acids or lipids also triggers SG formation^26, 27^. However, a combination of glycolytic and mitochondrial inhibition diminishes SG assembly^28^. Therefore, the exact effects of cellular energy metabolism on SG dynamics and the underlying mechanisms involved remain unclear.

The energy demand of the central nervous system (CNS) is high^29^, with neurons constantly consuming energy to maintain action potentials and synaptic function. Adenosine triphosphate (ATP) is the main currency of CNS energy metabolism, which is fueled by glucose catabolism through glycolysis and mitochondrial oxidative phosphorylation (OXPHOS)^30, 31^. Increasing evidence suggests that disrupted energy metabolism in CNS is involved in the development of ALS/FTD. Reduced glucose metabolism is observed in the motor-sensory cortex of ALS patients and in the frontal lobes, striatum, and thalamus of FTD patients, including in asymptomatic carriers of the ALS/FTD-linked C9orf72 mutation^32–34^. Glycogen, the storage form of glucose, is elevated in ALS mice and in patients’ spinal cord tissues^35, 36^. Moreover, mitochondrial dysfunction is a clinical hallmark of ALS/FTD and is mechanistically associated with several genetic forms of the disease, including those linked to SOD1, TDP-43, FUS, CHCHD10, and C9orf72^37–42^. A clinical study suggests that dietary intervention with high fat or high carbohydrate benefits ALS patients^43^. The mechanism by which disrupted energy metabolism contributes to the pathogenies of ALS/FTD, however, has not been fully elucidated.

The formation of conventional SGs is often triggered by phosphorylation of the translation initiation factor eIF2α, leading to polysome runoff and release of free mRNAs^3, 5^. Phosphorylated eIF2α (peIF2α) disrupts the translation-competent eIF2α-GTP-tRNA_i_^Met^ ternary complex, leading to stalled translation initiation. Alternatively, several small molecules can induce SG formation independently of peIF2α by disrupting the cap-binding eIF4F complex, which consists of eIF4E, eIF4A, and eIF4G^3^. For example, the eIF4A inhibitor hippuristanol binds to eIF4A, a DEAD-box helicase, and blocks its ATP-dependent RNA binding, thereby disrupting the function of the eIF4F complex and triggering the formation of an unconventional subtype of SG featuring RNA condensation occurring in a G3BP1/G3BP2-indepedent manner. In addition, selenite induces the formation of another unconventional subtype of SG, one which lacks eIF3, by dephosphorylating 4EBP1, leading to sequestration of eIF4E and therefore inhibition of eIF4F^44^. However, it remains unknown whether intracellular stress regulates SG formation through the eIF4F complex.

Here, we report that two energy-related metabolic pathways, glycolysis and mitochondrial OXPHOS, regulate the balance between the formation and clearance of SGs by maintaining the intracellular ATP pool. We found that blocking glycolysis, but not the pentose phosphate pathway (PPP), induced rapid formation of a new type of granule, as a direct consequence of a severe ATP reduction by more than 50%. Since these granules had a unique protein composition and exhibited different shapes and dynamics than those of the arsenite-induced SGs, we have termed them “energy-associated stress granules” (eSGs) and hereafter refer to the conventional arsenite-induced stress granules as “SGs”. Consistent with its role in glycolysis, cell-intrinsic glycogen proved to be critical for regulating eSG formation, when extracellular glucose concentration was within the physiological range. As the indispensable source of intracellular ATP when glycolysis is limited, mitochondrial OXPHOS was another critical regulator of eSG formation. Intracellular ATP also plays a vital role in regulating the clearance of conventional SGs; we found that moderate ATP reduction delayed SG clearance without triggering eSG formation. Mechanically, the ATP availability controls the formation of eSGs by regulating eIF4F assembly and RNA condensation, while the RNA condensation also underlies the ATP-regulated clearance of conventional SGs. An upregulation in eSG formation and reduction in SG clearance were observed in ALS patient-derived motor neurons and cortical organoids in response to energy stress, suggesting that these phenomena are part of the pathogenic mechanisms contributing to the neurodegenerative diseases.

## Results

### Glycolytic inhibition induces rapid eSG formation through a loss of ATP

Glycolysis is a major pathway of energy metabolism that breaks down glucose imported from the extracellular environment or glycogen stored intracellularly to generate pyruvate and ATP via multiple enzymatic steps (Fig. 1a). We first analyzed the influence of glycolysis on stress granule dynamics. To block the glycolysis pathway completely, we used glucose deprivation (GD) in combination with the administration of a specific glycogen phosphorylase (GP) inhibitor, CP91149 (CP), which blocks the breakdown of cell-intrinsic glycogen into glucose-1-phosphate (G1P) for subsequent catabolism, to shut down glucose and glycogen metabolism simultaneously (GD + CP) (Fig. 1a)^45^. Alternatively, we used 2-deoxy-D-glucose (2DG), a glucose analog that cannot be utilized by glycolysis and competitively inhibits hexokinase (HK) and glucose-6-phosphate isomerase (GPI), to inhibit glycolysis fueled by both glucose and glycogen (GD + 2DG) (Fig. 1a)^46, 47^. We found that robust cytoplasmic eSGs formed rapidly in HeLa cells stably expressing mCherry-G3BP1 after a 1-h inhibition of glycolysis (Fig. 1b). Like the conventional SGs induced by arsenite, eSGs contained multiple stress granule markers, including two RBPs, HUR and PABP, a translation initiation factor eIF4G, a 40S ribosomal subunit RPS6, and an ALS/FTD associated RBP TDP-43 (Fig. 1b and Extended Data Fig. 1a). Moreover, *in situ* hybridization showed strong recruitment of bulk mRNAs to eSGs (Fig. 1c). The eSGs were highly dynamic and exhibited liquid behaviors such as fusion to form larger granules (Extended Data Fig. 1b).

**Fig. 1.**
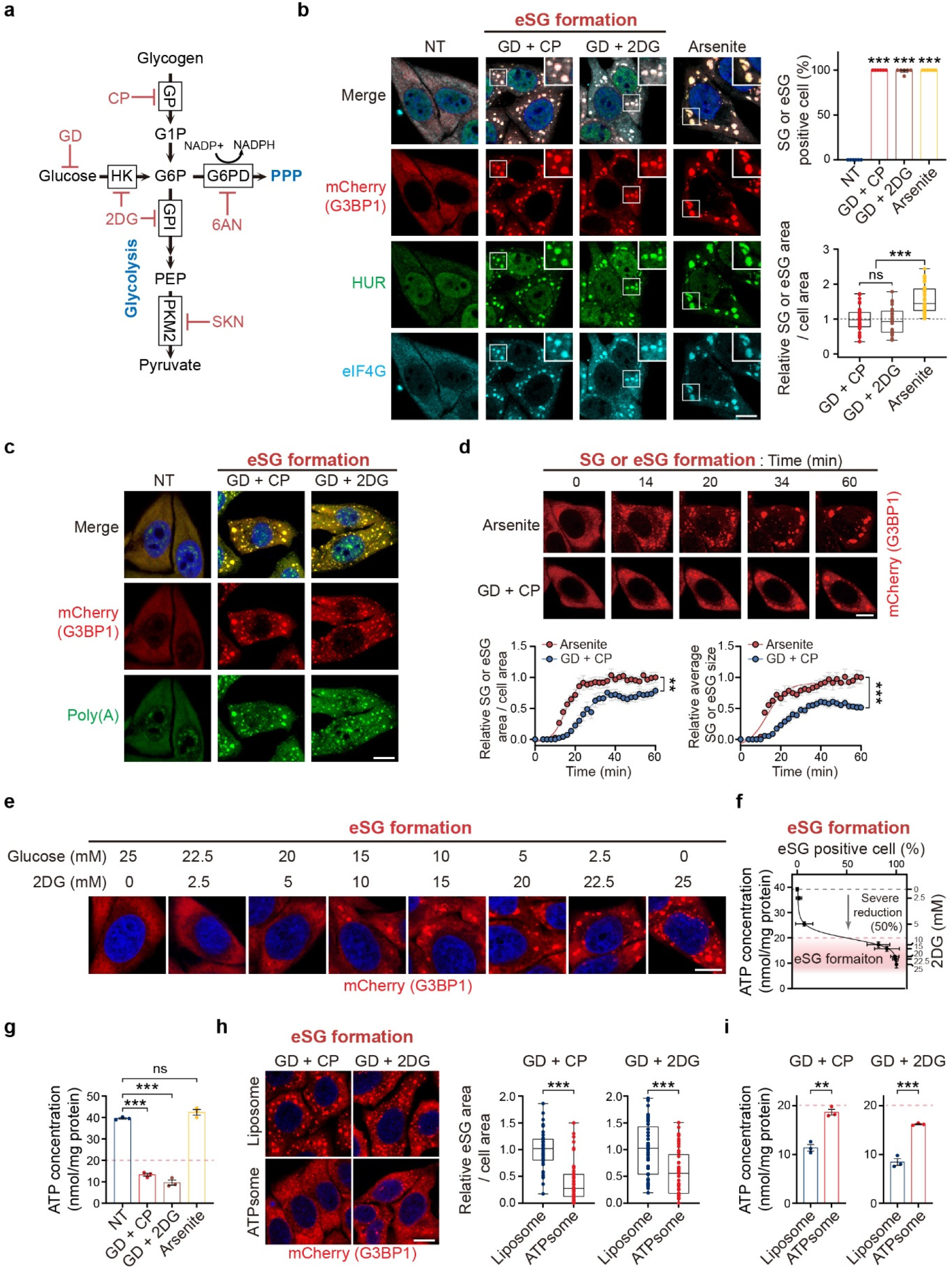
Severe ATP reduction by glycolytic inhibition prevents eSG formation. **a**, Schematic depiction of the glycolysis and pentose phosphate pathways (PPP) and targets of inhibitors (displayed in red) used in this study. **b**,**c**, Representative images of eSGs in HeLa cells stably expressing mCherry-G3BP1 and exposed to glucose deprivation (GD) in the presence of CP (50 μM) or 2DG (25 mM) for 1 h. Conventional SGs in cells treated with arsenite (250 μM) for 1 h served as controls. Cells were immunostained with anti-HUR and anti-eIF4G antibodies (**b**) or RNA fluorescence *in situ* hybridization (FISH) using an oligo-dT_20_ probe (**c**). The percentages of conventional SG- or eSG-positive cells (n = 6) or the relative areas of the total granules per cell (n = 35 cells) were assessed by mCherry-G3BP1 (**b**). NT: no treatment. **d**, Time-lapse imaging of SG or eSG formations in cells stably expressing mCherry-G3BP1 after treatment with arsenite or glycolysis inhibition for 1 h, respectively. The relative values of the granule area or average individual granule size per cell were quantified and normalized to the maximum value of SGs in the arsenite group (n = 10 cells from three independent experiments). **e**, Representative images of eSG formation in mCherry-G3BP1-expressing cells treated with increasing concentrations of 2DG along with decreasing glucose concentrations for 1 h. **f**, Intracellular ATP concentrations of cells treated as in (**e**) were measured (n = 3) and plotted against the eSG formation capacity in these cells as assessed by the percentage of eSG-positive cells (n = 6). The ATP concentration range correlating with robust eSG formation after a severe ATP reduction (50% reduction) is indicated. **g**, Intracellular ATP concentrations of cells undergoing the indicated treatments (n = 3). The ATP level below which eSG formation was triggered is indicated by the dotted line. **h**, Representative images and quantification of eSG formation, as assessed by the relative eSG areas per cell, in mCherry-G3BP1-expressing cells treated with the indicated energy stress for 1 h in the presence of either control empty liposomes (Liposome) or ATP-containing liposomes (ATPsome) (n = 45 cells from three independent experiments). **i**, Intracellular ATP concentrations in cells treated as in (**h**) (n = 3). The ATP level below which eSG formation was triggered is indicated. Nuclei were visualized by DAPI staining (blue). Data are shown as means ± SEM, analyzed by unpaired two-sided Student’s *t*-test. * P < 0.05; ** P < 0.01; *** P < 0.001; ns, not significant. Scale bars, 10 μm.

We next compared the assembly features of the glycolytic blockage-induced eSGs and of conventional arsenite-induced SGs. The total granule area of eSGs per cell was smaller than that of the conventional SGs (Fig. 1b). We further monitored the time course of granule assembly in cells treated with arsenite or exposed to glycolytic blockage. Using live-cell imaging, we found that the formation of arsenite-induced SGs started about 12 min after stress induction, and the area occupied by total or individual SG(s) continually increased to reach a maximum by 25 min after induction (Fig. 1d). In contrast, the assembly start time of eSGs was slightly but consistently delayed (∼16 min), and the growth of the total or individual eSG area took 40 min to reach its maximum, illustrating differing dynamics in the assembly of eSGs and arsenite-induced conventional SGs.

In addition to glycolysis, the pentose phosphate pathway (PPP) is an important branch of glucose and glycogen metabolism for producing NADPH and maintaining redox balance. To determine whether eSG formation induced by simultaneously blocking glucose and glycogen metabolism is independent of the PPP, we treated cells with 6-aminonicotinamide (6AN), a specific inhibitor of the PPP that acts by inhibiting G6PD, the first enzyme to shunt glucose and glycogen metabolism toward the PPP branch (Fig. 1a)^48^. For comparison, we also employed a specific glycolysis inhibitor, shikonin (SKN), which targets only the last glycolytic enzyme pyruvate kinase M2 (PKM2) and therefore does not affect PPP activity (Fig. 1a)^49^. Whereas robust eSG formation occurs in the presence of SKN, no granules were observed in cells treated with 6AN, indicating that inhibition of glycolysis, but not the PPP, had triggered eSG formation (Extended Data Fig. 1c).

Overexpression of G3BP1 is sufficient to trigger stress granules in the absence of stress^50^. To rule out the possibility that the eSGs were formed at least partially as a result of the exogenously expressed mCherry-G3BP1, we repeated the glycolytic blockage assays in normal HeLa cells and analyzed the eSG formation using immunostaining against endogenous G3BP1. Similar levels of eSG formations were observed (Extended Data Fig. 1d). Moreover, robust glycolytic blockage-induced eSGs were observed in mouse embryonic fibroblasts (MEF) and human retinal pigment epithelial (RPE1) cells (Figures S1E and S1F), further confirming that blockage of glycolysis was sufficient to induce eSG formation in mammalian cells.

Conventional SG formation is usually associated with translation arrest and disassembly of polysomes. We next asked whether eSGs also exhibit these two features. Ribopuromycinylation assay showed that blockage of glycolysis significantly reduced the global translation (Extended Data Fig. 1g). To determine whether eSG formation requires polysome disassembly, we treated cells with arsenite or inhibited the glycolysis in the presence of cycloheximide, a translation elongation inhibitor that leads to polysome stabilization. Cycloheximide strongly mitigated the formation of SGs or eSGs induced by arsenite or glycolytic inhibition, respectively, suggesting that eSG formation also depends on polysome disassembly (Extended Data Fig. 1h).

To gain molecular insights into the eSGs, we analyzed the protein composition of the eSGs and conventional SGs by mass spectrometry analyses after isolation of these granules (Extended Data Fig. 2a)^28, 51^. We identified 793 proteins that were enriched in either one or both granule fractions by more than 3-fold in the stressed cells as compared to the nontreated controls. Among these 793 proteins, 314 were identified in both granule fractions, whereas 257 and 222 proteins were enriched specifically in eSGs and conventional SGs, respectively (Extended Data Fig. 2b and Supplementary Table 1). Gene ontology (GO) pathway analysis indicated that the proteins identified in both eSGs and SGs were clustered into two top biological processes, RNA metabolism and translational initiation (Extended Data Fig. 2d). This result is consistent with the fact that stress granules are composed of RNPs and translation initiation factors, indicating that eSGs and SGs share common key stress granule components. We also found that the top 100 SG-specific proteins were enriched in the nucleocytoplasmic transport category; in contrast, the top 100 eSG-specific proteins were enriched in the mRNA maturation, nuclear DNA maintenance, and gene transcription categories (Extended Data Fig. 2d). Consistent with this observation, GO cellular compartment analysis indicated that more than half of the SG-specific proteins were cytosolic proteins; in contrast, most eSG-specific proteins were nuclear proteins (Extended Data Fig. 2e). The compartmentalization of top-ranked SG-specific proteins (EIF2A, ACP1, TNPO3, and NUP107) and eSG-specific proteins (NBN, hnRNPM, MRE11, and RMI1) were then confirmed by immunofluorescence staining (Extended Data Fig. 2f). Together, these data further confirm the shared nature of eSGs and SGs; however, the unique protein composition of the eSGs suggests they may have different functions than those of conventional SGs in the stress response.

Glycolysis is a major cellular energy source, and therefore, we asked whether the loss of ATP is the driver for eSG formation. Using 2DG, which competitively inhibits glucose and glycogen utilization, we gradually blocked cellular glycolytic activity by treating the cells with increasing concentrations of 2DG and decreasing concentrations of extracellular glucose and monitored their eSG formation and intracellular ATP levels (Fig. 1e). We found that the reduction in ATP levels was proportional to the increases in eSG formation, suggesting a strong correlation between ATP reduction and eSG formation (Fig. 1e,f). The eSGs began to appear when the ATP level was severely reduced by 50% and reached their maximum level when ATP was further reduced by 60% to 80% (Fig. 1f). We also observed comparable ATP reductions in cells with eSGs induced by the other glycolysis-blocking method (GD + CP) (Fig. 1g). In contrast, there was no change in the ATP level in cells with conventional SGs induced by arsenite treatment. To confirm that the severely reduced ATP level was the cause of the eSG formation when glycolysis was blocked, we boosted the ATP levels in cells with blocked glycolysis by administering ATP-containing liposomes (ATPsome) and found that the cells’ eSG formation was significantly reduced (Fig. 1h,i). Therefore, these data demonstrate that glycolytic inhibition can trigger the assembly of eSGs by causing severe energy stress when the ATP level is reduced by more than 50%.

### Cell-intrinsic glycogen is critical for preventing eSG formation at physiological levels of extracellular glucose

Glycolysis is fueled by both extracellular glucose and cell-intrinsic glycogen. Since the glucose concentration in the standard commercial culture medium is much higher than physiological concentrations, we asked how the eSG dynamics are controlled by glycolysis in the physiological range of extracellular glucose. In the brain, the glucose concentrations are 0.16 mM in hypoglycemia, 1 to 2.4 mM in normoglycemia, and 2.7 to 4.5 mM in hyperglycemia^52, 53^. Therefore, we tested the effects of physiologically high glucose (5 mM), normal glucose (1 mM), or low glucose (0.25 mM) on eSG formation. We were unable to induce acute eSG formation after a 1-h exposures to any of the three conditions (Fig. 2a,b). Interestingly, glucose deprivation alone did not induce eSG formation either (Fig. 2a,b). Given that robust eSGs were observed in cells treated with glucose deprivation plus blockage of glycogenolysis (Fig. 1a,b), these data collectively suggest that cell-intrinsic glycogen is crucial for preventing eSG formation when extracellular glucose is limited or deprived. Consistent with this concept, ATP concentrations in cells with intact glycogen metabolism could be maintained at a level that did not trigger eSG formation when extracellular glucose was reduced, accompanied by rapid consumption of intrinsic glycogen, suggesting that glycogen-fueled glycolysis is sufficient to sustain ATP levels and prevent eSG formation (Fig. 2c,d). Next, cells in each glucose condition were treated with CP to block glycogenolysis, as indicated by the stabilization of glycogen levels (Fig. 2d). We found that, when glycogenolysis was concomitantly blocked, cells treated with physiologically normal levels of glucose began to form mini-eSGs; furthermore, the eSG formation became more prominent under the low-glucose condition, comparable to that in cells with complete glycolytic blockage (GD + CP) (Fig. 2a,b). Consistently, the ATP levels were severely reduced by >50% in these cells exhibiting eSGs, further confirming that a half reduction in the ATP pool was the driver for the eSG formation (Fig. 2c).

**Fig. 2.**
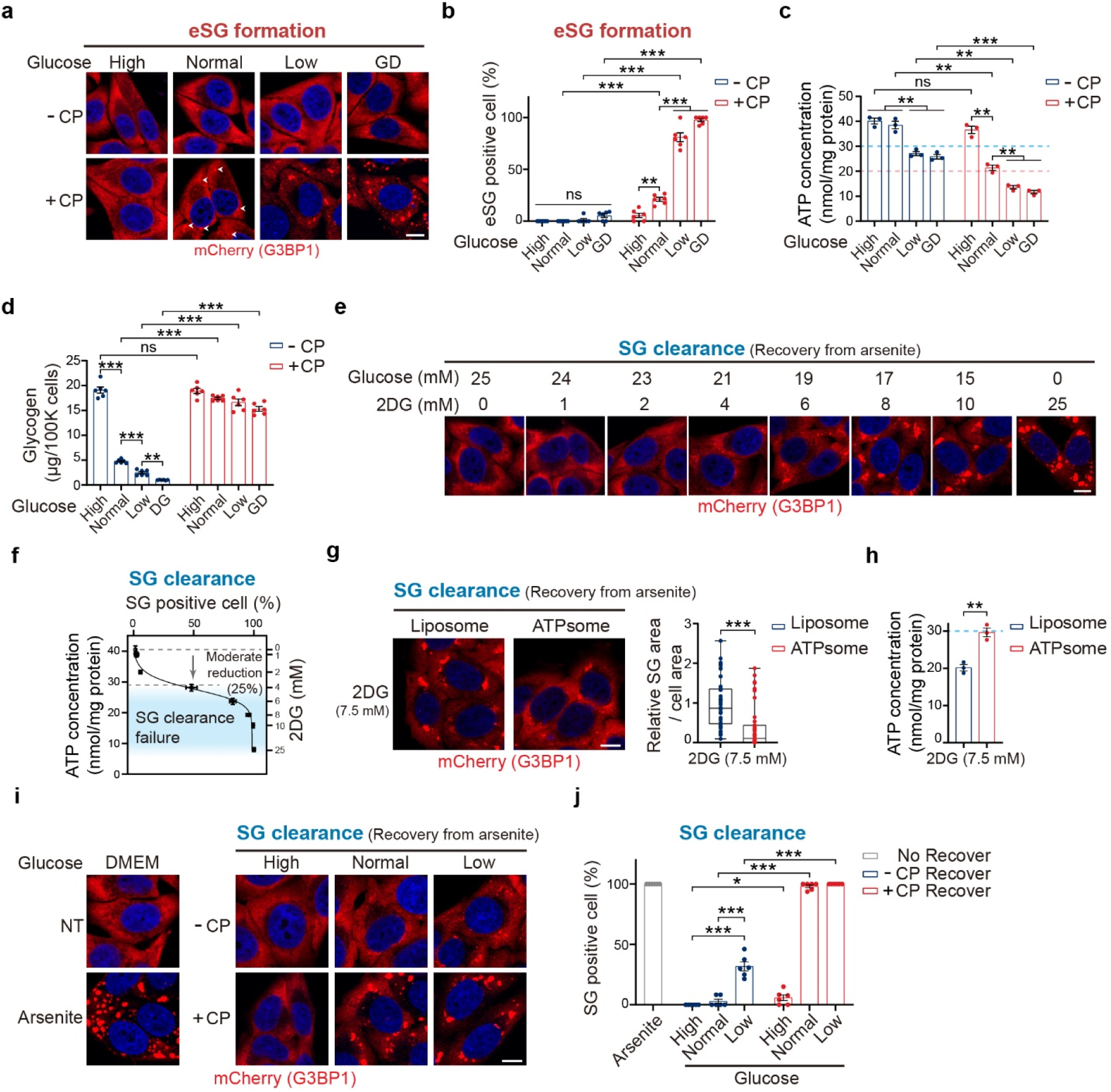
Cell-intrinsic glycogen and physiological levels of extracellular glucose fuel glycolysis to regulate eSG formation and maintain SG clearance. **a**, Representative images of eSGs in HeLa cells stably expressing mCherry-G3BP1 and treated with the indicated physiological range of glucose or by glucose deprivation (GD) for 1 h, with or without glycogenolysis inhibition produced by administration of CP. Arrowheads indicate the presence of mini eSGs. **b**, Quantification of eSG formation as assessed by the percentages of eSG-positive cells in (**a**) (n = 6). **c**,**d**, Intracellular ATP concentrations (**c**) (n = 3) or glycogen levels (**d**) (n = 6) in HeLa cells treated as in (Aa) (n = 3). The ATP levels below which eSG formation was triggered (pink) or SG clearance was impaired(cyan) are indicated by dotted lines (**c**). **e**, Representative images of persistent SGs in mCherry-G3BP1-expressing cells treated first with arsenite for 1 h in complete medium and then allowed to recover for 1 h after arsenite removal in medium containing increasing concentrations of 2DG and decreasing concentrations of glucose. **f**, Intracellular ATP concentrations in cells treated as in (**e**), measured (n = 3) and plotted against the levels of persistent SGs in these cells, as indicated by the percentage of SG-positive cells (n = 6). The ATP concentration range that was correlated with SG clearance failure after a moderate ATP reduction (by 25% to 80%) is highlighted. **g**, Representative images and quantification of persistent SGs, as assessed by the relative SG area per cell, in mCherry-G3BP1-expressing cells exposed to arsenite and then allowed to recover for 1 h after arsenite removal in medium containing 7.5 mM 2DG and 17.5 mM glucose plus either control empty liposomes or ATP-containing liposomes (ATPsome) (n = 45 cells). **h**, Intracellular ATP concentration in cells treated as in (**g**) (n = 3). The ATP level below which the SG clearance was impaired is indicated. **i**, Representative images of persistent SGs in mCherry-G3BP1-expressing cells that had recovered from arsenite treatment in medium with the indicated physiological range of glucose, with or without the inhibition of glycogenolysis by CP administration. **j**, Quantification of the SG clearance capacity, as assessed by the percentage of persistent SG-positive cells, in cells treated as in (**i**) (n = 6). Nuclei were visualized by DAPI staining (blue). Data are shown as means ± SEM, analyzed by unpaired two-sided Student’s *t*-test. * P < 0.05; ** P < 0.01; *** P < 0.001; ns, not significant. Scale bars, 10 μm.

To confirm that the critical function of glycogen in preventing eSG formation is to act as a fuel reserve for glycolysis under conditions of physiologically low glucose, we augmented the cells’ capacity to use environmental glucose by overexpressing the glucose transporter 1 (GLUT1), which is ubiquitously expressed in most cell types (Extended Data Fig. 3a). Overexpression of GLUT1 was sufficient to increase glucose uptake and partially restored cellular ATP concentrations to a level that prevented eSG formation in cells exposed to low glucose together with blockage of glycogenolysis (Extended Data Fig. 3b,c). We found that eSG formation was decreased in cells transfected with GLUT1, in contrast to the robust eSG observed in the non-transfected or GFP-transfected cells (Extended Data Fig. 3d,e), confirming that glycolysis of cell-intrinsic glycogen is crucial for the abilities of cells to sustain cellular ATP levels and prevent eSG formation at physiological levels of glucose.

### Glycolysis fuels the clearance of SGs

RNP granule dynamics are regulated by both assembly and clearance processes. We next examined whether either glycolysis or intracellular energy status also regulates the clearance of stress-induced granules. Since the formation of eSGs already cause a decrease in cellular ATP, it is infeasible to further analyze the effects of energy stress on the process of eSG clearance. Unlike eSGs whose induction depends on energy deprivation, the formation of conventional SGs induced by arsenite does not involve a disturbance in the cell’s energy status (Fig.1 g), allowing us to test the effects of energy stress on granule clearance. Specifically, cells were first stressed by arsenite treatment under normal culture conditions and then allowed to recover in various energy stress media without arsenite. We first inhibited glycolysis in the cells undergoing SG clearance by gradually increasing the cellular concentrations of 2DG while decreasing the glucose concentrations. We found that blocking glycolysis significantly delayed SG clearance (Fig. 2e). Moreover, by determining the energy status of these cells, we noted a strong correlation between SG clearance rates and intracellular ATP concentrations, with the SG clearance rate beginning to drop when the intracellular ATP level was moderately reduced by >25% (Fig. 2f). Administration of ATP-containing liposomes restored the ATP level and the SG clearance capacity, suggesting that the moderately reduced ATP level was the cause of the delayed SG clearance (Fig. 2g,h). Thus, these data indicate that adequate glycolytic activity and ATP supply are required for effective SG clearance.

Next, we analyzed the effects of glycogen metabolism and the physiological range of extracellular glucose on SG clearance. We found that cells under high or normal glucose conditions could clear SGs in 1 h after arsenite removal (Fig. 2i,j). In comparison, about 30% of cells under low-glucose condition still harbored visible SGs after 1 h, suggesting that a physiologically normal level of extracellular glucose is required for effective SG clearance. We further tested the role of glycogen on SG clearance but interestingly found no SG clearance in cells under normal or low-glucose conditions when glycogenolysis was blocked (Fig. 2i). Given that the ATP concentrations in the cells with impaired SG clearance were reduced by more than 25% (Fig. 2c), this result suggests that endogenous stored glycogen had served as an indispensable fuel source for glycolysis to produce ATP when extracellular glucose was limited and that the depletion of glycogen in cells under low-glucose conditions led to moderate energy stress that impaired SG clearance. To further evaluate this idea, we examined the SG clearance capacity of cells whose glucose uptake was enhanced through elevation of GLUT1 expression. We found that GLUT1 overexpression significantly accelerated the SG clearance rates in cells when glycogenolysis was blocked under the condition of normal glucose, consistent with the restoration of ATP in these cells (Extended Data Fig. 3f-h), and confirming that glycogen regulates the SG clearance by acting as an alternative fuel source for glycolysis.

### OXPHOS influences eSG formation and SG clearance through ATP

In addition to glycolysis, mitochondrial OXPHOS serves as another major cellular energy source by producing ATP through acetyl-CoA oxidation. Therefore, we next tested the role of OXPHOS in regulating eSG formation. It should be noted that cells have evolved to shift quickly from OXPHOS to glycolysis to sustain the energy supply when OXPHOS is inhibited. Therefore, it is crucial to analyze the effects of OXPHOS in cells fed with glucose at concentrations within the physiological range. No eSGs were observed in cells treated with oligomycin, a chemical blocking the ATP synthase of OXPHOS, under conditions of physiologically high or normal glucose (Fig. 3a,b). Consistent with this observation, ATP levels in the oligomycin-treated cells were maintained at above 50%, a level that does not trigger eSG formation (Fig. 3c). These data suggest that high or normal glucose levels can enable cells to produce sufficient ATP to prevent eSG formation via glycolysis when OXPHOS is blocked. However, ATP levels in oligomycin-treated cells with the low glucose level were dramatically decreased (Fig. 3c). Consistently, robust eSG formation was observed in the cells under this severe energy-stress condition (Fig. 3a,b). As was seen for the eSGs induced by blocking glycolysis, the eSGs triggered by OXPHOS inhibition contained multiple stress granule markers and bulk mRNA, and their formation required polysome disassembly (Extended Data Fig. 4a-c). Moreover, partially restoring ATP concentrations by administering ATP-containing liposomes decreased the eSG assembly (Extended Data Fig. 4d,e). Together, these data indicate that ATP produced by OXPHOS is indispensable for supplying cellular energy and preventing eSG formation when environmental glucose is low.

**Fig. 3.**
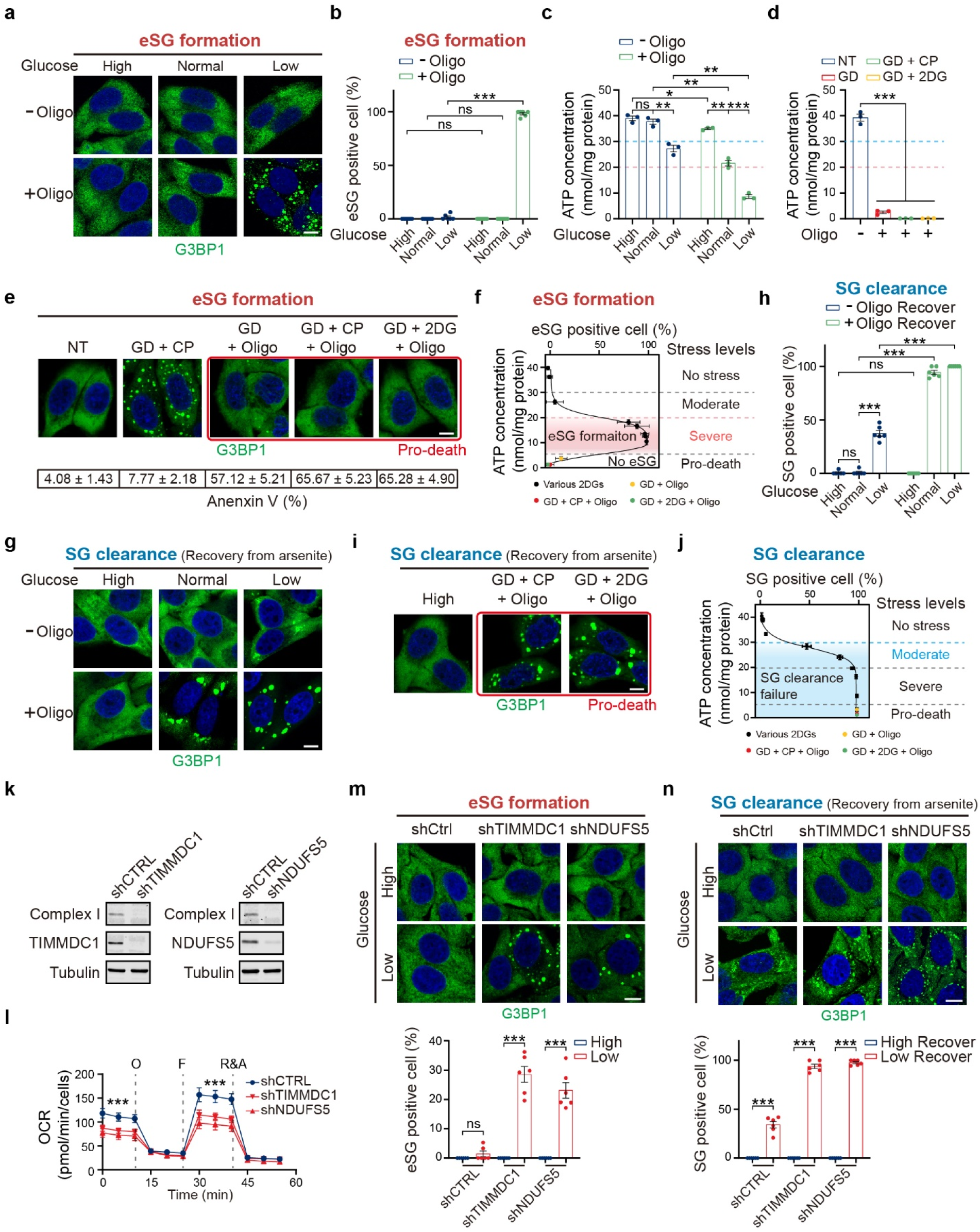
OXPHOS is critical for preventing eSG formation and sustaining SG clearance in physiological levels of glucose. **a**,**b**, Representative images (**a**) and quantification (**b**) of eSG formation, visualized by G3BP1 immunofluorescence analysis (IF) and assessed by the percentage of eSG-positive cells (n = 6) in HeLa cells treated with the indicated physiological range of glucose for 1 h, with or without OXPHOS inhibition by oligomycin (Oligo, 2 μM). **c**,**d**, The intracellular ATP concentrations in cells treated as indicated for 1 h (n = 3). The ATP levels below which eSG formation was triggered (pink) or SG clearance was impaired (cyan) are indicated by dotted lines. **e**, eSG formation as assessed by G3BP1 IF in cells exposed to 1 h of either glycolysis inhibition that triggered eSG formation (GD + CP) or the indicated pro-death energy stress highlighted in red. Cell death (%) in each group was examined by Annexin V staining and is shown beneath (n = 4). **f**, Intracellular ATP concentrations in cells treated with the indicated pro-death energy stress as in (**e**) (n = 3), plotted against eSG formation capacity in these cells, as assessed by the percentage of eSG-positive cells (n = 6). The various levels of energy stress and the ATP range that triggered eSG formation are marked for the data shown here, in combination with the data for cells treated with increasing concentrations of 2DG in (Fig. 1f). **g**, Representative images of persistent SGs in HeLa cells after recovery from arsenite treatment for 1 h in the indicated physiological glucose range, with or without inhibition of OXPHOS with oligomycin. SGs were visualized by G3BP1 IF. **h**, Quantification of SG clearance capacities, assessed by the percentage of SG-positive cells, in the cells from (**g**) (n = 6). **i**, Persistent SGs in HeLa cells after recovery from arsenite treatment in the indicated pro-death energy stresses for 1 h, as assessed by G3BP1 IF. **j**, Intracellular ATP concentrations of cells treated with indicated pro-death energy stress as in (**i**) were measured (n = 3) and plotted against the level of persistent SGs in these cells, as assessed by the percentages of SG-positive cells (n = 6). The ATP range that impaired the SG clearance when the energy stress level was moderate or higher is marked for the data shown here, in combination with the data for cells treated with increasing concentrations of 2DG in (Fig. 2f). **k**,**l**, Mitochondrial complex I levels (**k**) and oxygen consumption rates (OCRs) (**l**) (n=6) in HeLa cells stably transduced with a TIMMDC1 or NDUFS5 shRNA. The efficiency of knockdown of TIMMDC1 or NDUFS5 was confirmed by immunoblotting. Complex I levels were determined by immunoblotting against one of its subunits, NDUFB8 (**k**). shCTRL: non-targeting control shRNA. O, oligomycin; F, FCCP; R&A, rotenone and antimycin A. **m**,**n**, Representative images of eSGs (**m**) or persistent SGs after recovery from arsenite treatment (**n**) in control cells or cells with knockdown of TIMMDC1 or NDUFS5 fed with the indicated glucose concentrations for 1 h. The percentages of eSG- or SG-positive cells were quantified (n = 6). Nuclei were visualized by DAPI staining (blue). Data are shown as means ± SEM, analyzed by unpaired two-sided Student’s *t*-test. * P < 0.05; ** P < 0.01; *** P < 0.001; ns, not significant. Scale bars, 10 μm.

We next examined the effects of more severe energy stress on eSG formation by simultaneously shutting down glycolysis and OXPHOS. Unlike the above-discussed severe energy stresses that reduced the intracellular ATP levels by no more than 80% by inhibiting either glycolysis or OXPHOS when the glucose concentration was physiologically low, we observed that simultaneous inhibition of glycolysis and OXPHOS dramatically reduced the intracellular ATP levels by > 95% (Fig. 3d). Such near-complete energy depletion led to rapid cell death (Fig. 3e). In addition, we found that these pro-death energy stresses did not trigger eSG formation (Fig. 3e,f). These data suggest that the formation of eSGs is an energy consuming process and that pro-death energy stresses trigger cell death instead of eSG formation because of the complete depletion of energy.

We next analyzed the effect of OXPHOS on conventional SG clearance. SGs were fully cleared in cells under the high-glucose condition combined with OXPHOS inhibition, in line with the unchanged ATP level in these cells, suggesting that cells can rely solely on glycolysis for energy production as well as SG clearance when environmental glucose is high (Fig. 3c,g,h). In contrast, we observed that OXPHOS inhibition completely abolished the SG clearance under either normal- or low-glucose conditions, in line with the moderately reduced ATP levels in these cells (Fig. 3c,g,h). Moreover, pro-death energy stresses also blocked the SG clearance process (Fig. 3i,j). Partially restoring ATP by administration of ATP-containing liposomes accelerated SG clearance (Extended Data Fig. 4f,g). These data indicate that because of the high sensitivity of the SG clearance process to ATP deficiency, OXPHOS activity is crucial for avoiding moderate cellular energy stresses and sustaining effective SG clearance when the environmental glucose is low or even normal.

To further confirm the effects of OXPHOS on eSG formation and SG clearance, we generated two cell lines, each expressing an shRNA targeting either a subunit (NDUFS5) or a key assembly factor (TIMMDC1) of complex I, the first enzymatic complex of the OXPHOS pathway. Knockdown of each of the two proteins led to a depletion of complex I and a significant reduction in the OXPHOS activity (Fig. 3k,l). When cultured under the low-glucose condition, about 30% of both types of OXPHOS-deficient cells showed eSG formation, in contrast to the non-targeting control cells in which no eSG was observed (Fig. 3m). Moreover, after arsenite treatment and then removal, the clearance of SGs in both types of OXPHOS-deficient cells fed with low glucose was less efficient than that observed in control cells (Fig. 3n).

Glutamine is the most abundant amino acid in the human body. In culture, glutamine is converted by the cells into alpha-ketoglutarate (αKG), which is a key metabolite in the mitochondrial trichloroacetic acid (TCA) cycle that is subsequently oxidized for ATP production through OXPHOS^54, 55^. We therefore tested the effects of glutamine-dependent OXPHOS activity on eSG formation and SG clearance by using glutamine deprivation. In cells grown in the low-glucose medium, glutamine deprivation triggered eSG formation in half of the cells and reduced the ATP concentration by about 50% (Extended Data Fig. 4h,i). Such eSG-triggering effects of glutamine deprivation could be mitigated by administering of dimethyl α-ketoglutarate (Dm-αKG), a cell-permeable form of αKG, suggesting that conversion of glutamine to αKG for OXPHOS and subsequent ATP production underlies the eSG-triggering effect of glutamine deprivation (Extended Data Fig. 4j). Furthermore, when SG clearance rates were analyzed, glutamine deprivation also significantly delayed this process in cells grown in normal or low-glucose medium (Extended Data Fig. 4k). Together, these data support the critical role of OXPHOS in preventing eSG formation and maintaining SG clearance when extracellular glucose is limited.

### Energy stress triggers eSG formation via the mTOR-4EBP1 axis

We next analyzed the molecular basis of eSG formation. We first tested the role of eIF2α phosphorylation (peIF2α), which is the most-studied molecular trigger for translation inhibition and conventional SG formation in response to various stressors. As a positive control, arsenite treatment caused a dramatic increase in the peIF2α level (Fig. 4a). However, under energy stress conditions that induced eSG formation, peIF2α levels were unchanged when compared to the nontreated control. Moreover, when the phosphorylation site (serine 51) in eIF2α was mutated (S51A), arsenite failed to trigger conventional SG formation (Fig. 4b); in contrast, the capacity for eSG formation in the mutant cells was similar to that in wild-type cells. These data suggest that eSG formation is triggered in a peIF2α-independent manner.

**Fig. 4.**
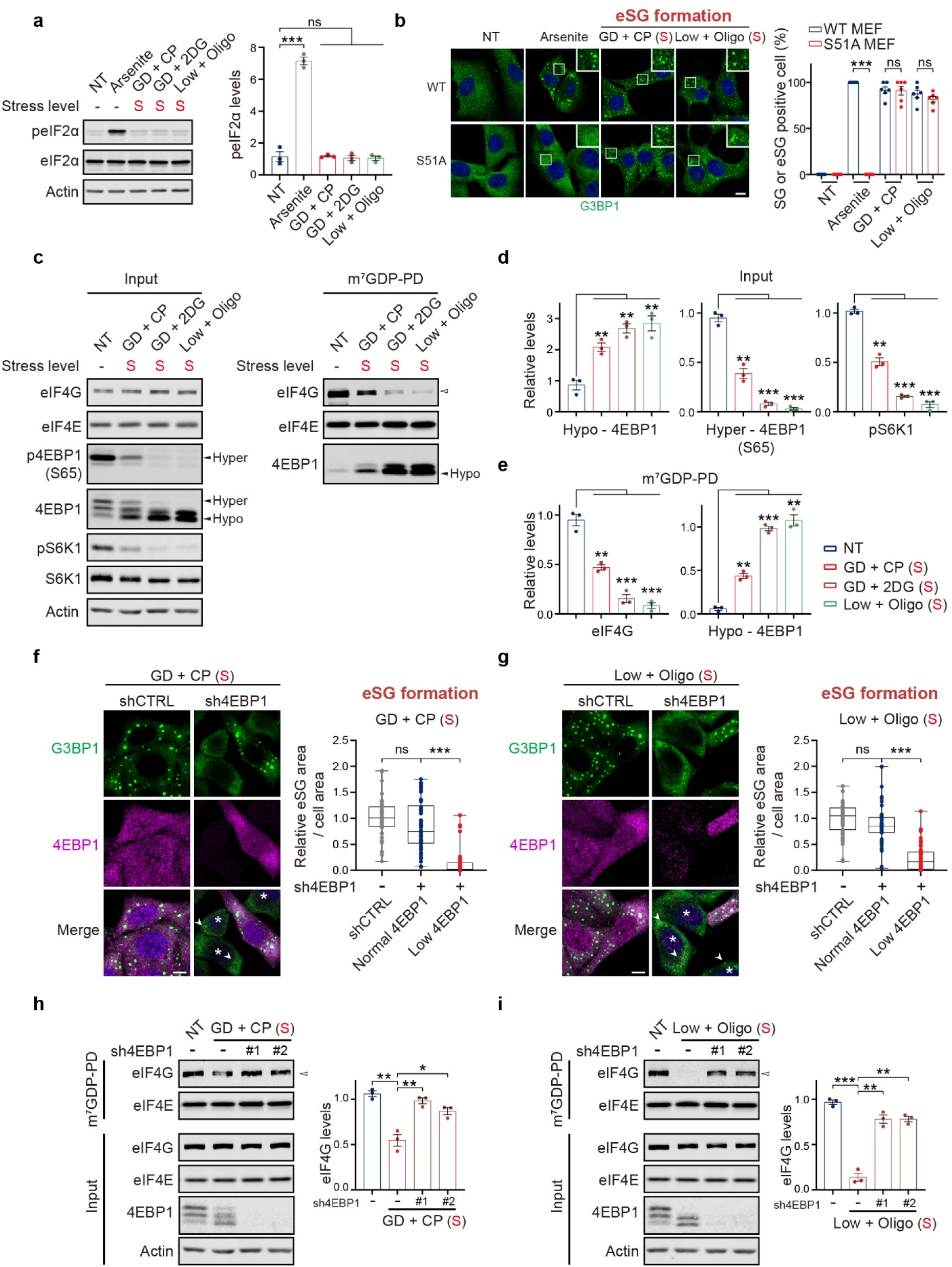
Severe energy stresses trigger eSG formation through 4EBP1 dephosphorylation. **a**, Western blot analysis of eIF2α phosphorylation (peIF2α) and total eIF2α levels in cells treated with the indicated severe energy stress or arsenite (n = 3). **b**, Wild-type MEFs (WT) or MEFs bearing S51A mutant eIF2α (S51A) were exposed to medium containing arsenite or the indicated severe energy stress for 1 h. Representative images of eSGs or SGs visualized by G3BP1 IF are shown. The percentage of eSG- or SG-positive cells was quantified (n = 6). **c**, HeLa cells exposed to one of the indicated severe energy stresses were lysed and subjected to m^7^GTP-Sepharose pulldown (PD) to isolate cap-associated proteins. Input lysates and pulldowns were analyzed by western blotting using the indicated antibodies. The positions of hyperphosphorylated (hyper) and hypophosphorylated (hypo) forms of 4EBP1 are indicated. The open arrowhead indicates the decreased association between eIF4G and m^7^GTP-Sepharose in stressed cells. **d**, Quantifications of the relative levels of hypo-4EBP1 as determined by using the total 4EBP1 antibody, hyper-4EBP1 levels as determined by Ser65 phosphorylation (S65), and pS6K1 levels in the inputs in (**c**) (n = 3). **e**, Quantifications of the cap-bound eIF4G levels and hypo-4EBP1 levels in the pulldowns in (**c**) (n = 3). **f**,**g**, HeLa cells were transfected with an shRNA-targeting 4EBP1 or a non-targeting control for 72 h and then subjected to one of the indicated severe energy stress treatments for 1 h. The eSGs in cells from the control group or in cells from the shRNA group with normal or low 4EBP1 levels were visualized by G3BP1 IF and quantified as the relative eSG area per cell (n = 50 cells from three independent experiments). Asterisks indicate cells with reduced 4EBP1 and diminished eSGs. Arrowheads indicate the presence of mini-eSGs in the 4EBP1-depleted cells. **h**,**i**, Western blot analysis of the association between eIF4G and the mRNA cap using the indicated antibodies. Cells were transfected with one of two shRNAs targeting 4EBP1 or a non-targeting control for 72 h and then subjected to m^7^GTP-Sepharose pulldown. Cap-associated eIF4Gs, indicated by open arrowheads, were quantified (n = 3). NT: no treatment. Nuclei were visualized by DAPI staining (blue). The stress levels of each treatment are indicated. S, severe energy stress. Data are shown as means ± SEM, analyzed by unpaired two-sided Student’s *t*-test. * P < 0.05; ** P < 0.01; *** P < 0.001; ns, not significant. Scale bars, 10 μm.

Cells respond to energy stresses via energy-sensing pathways. Adenosine monophosphate-activated protein kinase (AMPK) and mechanistic target of rapamycin (mTOR) are two core sensors that monitor cellular energy status and regulate various downstream effector pathways^56–58^. We next tested whether eSG formation was tied to these two energy-sensing pathways. AMPK is phosphorylated and activated by an increased AMP-to-ATP ratio when ATP consumption is elevated under low-energy conditions. We found that eSG-inducing severe energy stresses activated the AMPK pathway, as evidenced by increased AMPK phosphorylation levels (Extended Data Fig. 5a). Consistently, the phosphorylation of two well-established substrates of activated AMPK, acetyl-CoA carboxylase (ACC) and regulatory-associated protein of mTOR (raptor), was significantly upregulated. We next generated two independent HeLa cell lines with defective AMPK activity by disrupting both catalytic subunits (α1/α2) of AMPK (double-knockout, DKO) (Extended Data Fig. 5b). Notably, the eSG formation capacity in the two DKO cells was not affected by the loss of AMPK and was similar to that observed in wild-type cells, suggesting that eSG formation is independent of AMPK activity (Extended Data Fig. 5c).

mTOR also senses intracellular energy status and becomes inactivated when glycolysis or OXPHOS is inhibited. We found that the phosphorylation level of S6K1 at Thr389 (pS6K1) or the hyperphosphorylation level of 4EBP1 (indicated by Ser65 phosphorylation [p4EBP1]), two well-studied downstream targets of mTOR, was significantly decreased in cells exposed to severe eSG-inducing energy stresses (Fig. 4c,d). Moreover, the dephosphorylation of p4EBP1 resulted in a band shift toward the fastest-migrating band, the hypophosphorylated form of 4EBP1 (Fig. 4c). To further confirm that the dephosphorylation of 4EBP1 resulted from mTOR inactivation, we utilized two constitutively activated forms of mTOR (S2215Y and R2505P) to mimic its activation^59^. We found that overexpression of either of the two constitutively activated forms of mTOR restored the p4EBP1 level and decreased the faster-migrating, hypophosphorylated 4EBP1 in stressed cells (Extended Data Fig. 5d). These data indicate that the mTOR pathway is inactivated when eSG is triggered by severe energy stresses, leading to dephosphorylation of 4EBP1.

Upon dephosphorylation, 4EBP1 becomes active and then binds the translation factor eIF4E, which recognizes the 5’ cap of mRNA and recruits eIF4G, an essential step in eIF4F complex assembly and subsequent initiation of translation. The binding to 4EBP1 competitively prevents the interaction of eIF4E with eIF4G and represses translation. After pulling down the mRNA cap-binding proteins with m^7^GTP-Sepharose, we found that severe energy stress significantly decreased the levels of cap-bound eIF4G, and there was a concomitant increase in the interaction between 4EBP1 and cap-bound eIF4E (Fig. 4c,e). These data indicate that the dephosphorylation of 4EBP1 impairs the recruitment of eIF4G by cap-bound eIF4E and subsequent eIF4F assembly. Given that sodium selenite is known to cause translation inhibition and trigger an unconventional type of stress granule by inducing 4EBP1 dephosphorylation^44^, we hypothesized that severe energy stress blocks the initiation of protein translation and triggers eSG formation through 4EBP1 dephosphorylation resulting from mTOR inactivation. To test this possibility, we examined eSG formation in cells transduced with an shRNA targeting 4EBP1. We found that eSGs induced by two different severe energy stresses were dramatically diminished in cells showing reduced 4EBP1 levels when compared to those with normal levels of 4EBP1 or expressing a non-targeting control (Fig. 4f,g). Consistently, the reduction in recruitment of eIF4G to the mRNA 5’ cap when eSGs were triggered was corrected in 4EBP1-depleted cells (Fig. 4h,i). These data suggest that 4EBP1 dephosphorylation as a result of mTOR inactivation plays a critical role in eSG formation.

We also noted, however, that although the number of typical eSGs was largely decreased in the 4EBP1-depleted cells, a small number of visible mini-granules was still present in these cells (Fig. 4f,g). Whereas the granules induced by the selenite-induced dephosphorylation of 4EBP1 were characterized by the exclusion of eIF3^44^, we found that the eSGs did contain eIF3 components, as evidenced by IF staining of eIF3B and the detection of 12 out of 13 eIF3 proteins in a proteomic analysis of eSG components, suggesting that the composition of eSGs differs from that of the selenite-induced granules (Extended Data Fig. 5e,f and Supplementary Table 1). Moreover, forced dephosphorylation of 4EBP1, as a result of the pharmacological inactivation of mTOR by Torin1, did not induce eSG formation (Extended Data Fig. 5g,h). No ATP reduction was observed in these cells, consistent with the concept that a reduction in ATP is a key prerequisite for eSG formation (Extended Data Fig. 5i). In addition, although 4EBP1 knockdown was sufficient to restore the normal recruitment of eIF4G to the 5’ mRNA cap, global translation remained stalled (Fig. 4g,h and Extended Data Fig. 5j). Taken together, these data suggest that 4EBP1 dephosphorylation is an essential step in eSG formation; however, 4EBP1 dephosphorylation alone was insufficient to trigger eSG formation, and additional ATP-dependent molecular machinery may also be required for eSG formation in response to severe energy stresses.

### eIF4A is involved in energy-regulated eSG formation

DEAD-box ATPase proteins are RNA helicases that control RNA metabolism. As part of our search for additional players required for eSG formation, we focused on eIF4A, which is the most abundant DEAD-box protein in HeLa cells^60^. eIF4A prevents RNA condensation via its RNA binding capacity in an ATP-dependent manner, and pharmacologically interfering with eIF4A-RNA binding induces RNA condensation and the formation of unconventional stress granules independent of peIF2α^60, 61^. Therefore, we hypothesized that reducing ATP in cells exposed to various energy stresses impairs the ATP-dependent RNA-binding capacity of eIF4A1 and leads to eSG formation. To test this hypothesis, we first analyzed the RNA-binding capacity of eIF4A1 in cells exposed to severe energy stresses. By examining the association of eIF4A1 with its specific RNA targets by UV-crosslinking RNA immunoprecipitation (UV-RIP), we found that eSG-inducing severe energy stresses significantly decreased the association of eIF4A1 with its RNA targets, the *TFRC*, *DYNC1CH1*, and *AHNAK* mRNAs and the *NORAD* lncRNA, indicating that the RNA binding capacity of eIF4A1 is reduced when the intracellular ATP level is decreased (Fig. 5a). Moreover, eIF4A is known to prevent RNA condensation and stress granule formation by limiting the recruitment of bulk mRNA and the scaffold RBP G3BP1^60^. We found that significantly higher amounts of bulk mRNA and G3BP1 were partitioned into eSGs than into arsenite-induced conventional SGs, suggestive of relatively condensed RNAs in the eSGs (Fig. 5b,c). It should be noted that arsenite-induced SG formation requires the scaffold RBP of the multivalent network, G3BP1/G3BP2, to promote LLPS; however, RNA condensation, occurring as a result of the pharmacological inhibition of eIF4A can induce granule formation independently of G3BP1/G3BP2 by promoting multivalent RNA-RNA interactions^4, 60, 62^. Interestingly, we found that eSG formation still required the presence of G3BP proteins, suggesting that severe energy stress-induced RNA condensation is not as strong as that induced by the pharmacological inhibition of eIF4A1 and cannot induce LLPS independently of G3BP (Extended Data Fig. 6a). Together, these data indicate that, as a result of a reduction in ATP supply, eIF4A1 is less efficient in binding its RNA targets to prevent their condensations, together with the activated 4EBP1 function, leading to G3BP1 buildup and eSG formation.

**Fig. 5.**
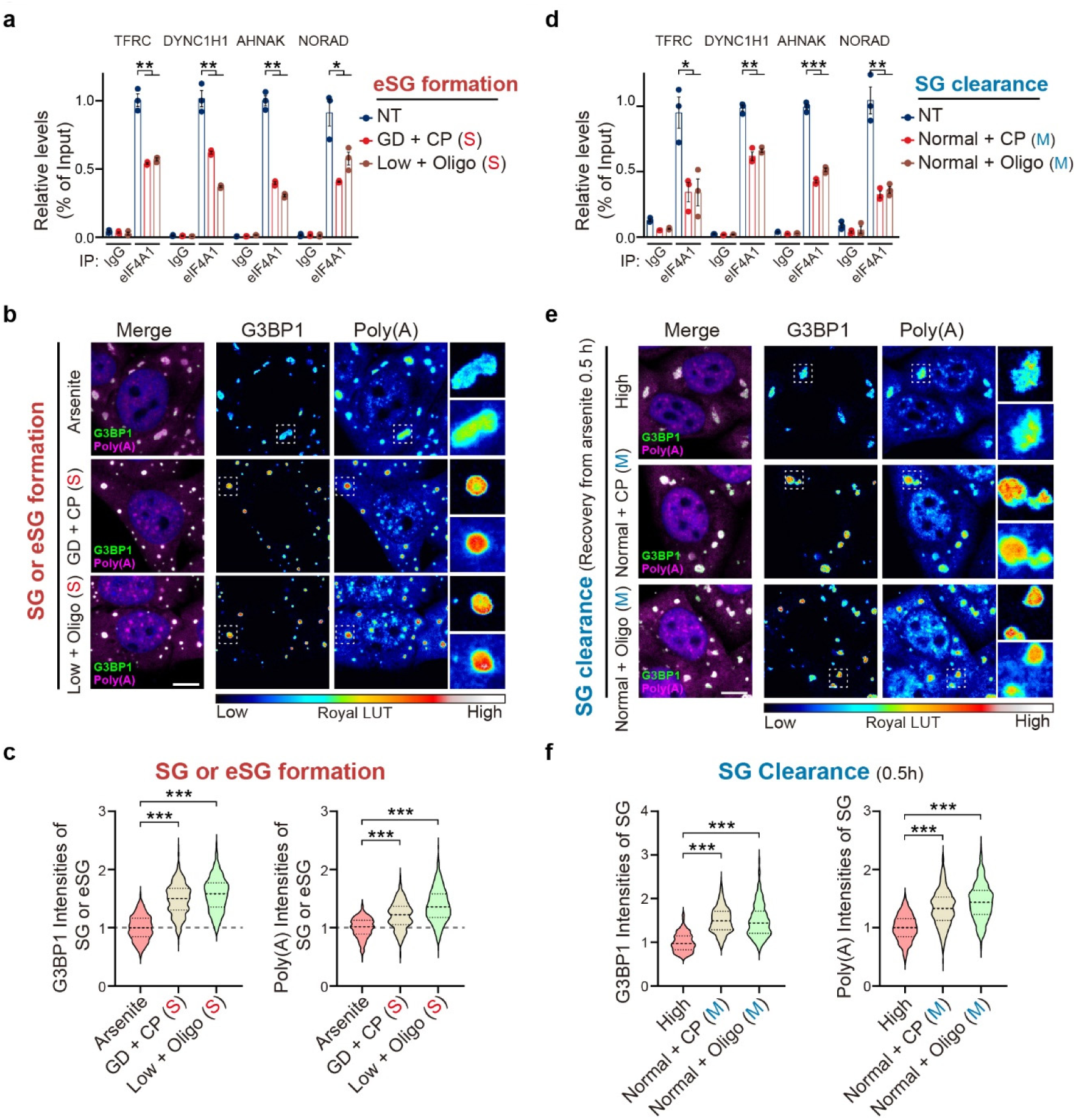
The reduced association of eIF4A with RNA is involved in energy stress-controlled eSG formation and SG clearance. **a**, UV-RIP of eIF4A1 was performed in HeLa cells treated with severe (S) energy stresses for 1 h, followed by qPCR to detect the enrichment of the *TFRC*, *DYNC1H1*, and *AHNAK* mRNAs, and the *NORAD* lncRNA. The level of each transcript is expressed as a percentage of the input (n = 3). **b**,**c**, Representative images of G3BP1 IF and mRNA FISH using an oligo-dT_20_ probe in HeLa cells treated with indicated severe energy stress (**b**). Images were captured at the same scale, and the fluorescent intensities of G3BP1 and mRNA are expressed as pseudocolor images. Magnified panels show the data for individual granules. Quantification of the intensity of G3BP1 or mRNA per granule is shown (**c**) (n > 30 cells from three independent experiments) **d**, UV-RIP of eIF4A1 was performed in HeLa cells after recovery from arsenite treatment in the indicated moderate energy stress for 1 h, followed by qPCR to detect the enrichment of the RNA targets as described in (**b**) (n = 3). **e**,**f**, Representative images of G3BP1 IF and mRNA FISH in HeLa cells after recovery from arsenite treatment in the indicated moderate energy stresses for 0.5 h (**e**). Pseudocolor images reflect the intensities of the indicated targets. Quantification of the intensity of G3BP1 or mRNA per granule was performed as described in (**f**) (n > 30 cells from three independent experiments). Nuclei were visualized by DAPI staining (blue). The stress levels of each treatment is indicated. S, severe energy stress; M, moderate energy stress. Data are shown as means ± SEM, analyzed by unpaired two-sided Student’s *t*-test. * P < 0.05; ** P < 0.01; *** P < 0.001; ns, not significant. Scale bars, 10 μm.

### eIF4A underpins energy-regulated SG clearance

We next explored the molecular mechanism underlying the process of clearing moderate energy stress-modulated conventional SGs. Since dephosphorylation of peIF2α is critical for the clearance of arsenite-induced conventional SGs after arsenite removal, we first assessed the peIF2α levels during the SG clearance in cells exposed to moderate energy stresses. Immunoblotting showed no difference in peIF2α dephosphorylation rates between cells under high glucose and those under various moderate energy stress conditions, suggesting that the delayed SG clearance induced by moderate energy stress does not involve the dephosphorylation of peIF2α (Extended Data Fig. 6b).

Since a moderate ATP reduction was found to be required for the delayed SG clearance process in response to moderate energy stresses, we next asked whether the two abovementioned ATP-sensitive regulators involved in eSG formation, namely 4EBP1 and eIF4A1, also played a role in the conventional SG clearance process. For 4EBP1, there was no difference in the levels of the translation-inhibiting, hypophosphorylated 4EBP1 in cells with or without exposure to moderate-energy stresses (Extended Data Fig. 6b), suggesting that, in contrast to the severe energy stresses triggering eSG formation, a moderate reduction in the intracellular ATP concentration does not disturb the 4EBP1 phosphorylation status. Therefore, the mTOR-4EBP1 pathway is not involved in regulating the clearance of eSGs modulated by moderate energy stresses.

Next, we tested whether the ATP-dependent RNA-binding capacity of eIF4A1 was affected by a moderate reduction in ATP. In the UV-RIP assay, we observed that the interactions of eIF4A1 with its specific RNA targets were significantly reduced in cells with moderate energy stresses (Fig. 5d). Moreover, when SGs were examined in the middle of the clearance process after arsenite removal for only 30 min, both bulk mRNA and G3BP1 were significantly enriched in the SGs in cells that has been exposed to moderate energy stresses, suggesting that the condensation of mRNA and G3BP1 is related to the impairment of eIF4A1 function in these cells (Fig. 5e,f). Therefore, our data suggest that a moderate reduction in intracellular ATP delays conventional SG clearance by disrupting the ATP-dependent RNA-binding capacity of eIF4A.

### Impaired eSG and SG dynamics in C9-ALS patients’ cells

Abnormal dynamics of stress-induced granules as a result of impaired energy metabolism have been implicated in the pathogenesis of NDDs, including ALS/FTD. We next asked whether severe energy stress-associated eSG formation and moderate energy stress-associated SG clearance were disrupted in patients’ cells. Considering that the *GGGGCC* repeat expansion in a noncoding region of the C9orf72 gene is the most common genetic cause of ALS/FTD and directly causes disrupted energy metabolism, especially OXPHOS, in patient-derived cells^39, 63^, we analyzed eSG formation under severe energy stress in C9-ALS patients’ iPSC-derived cortical organoids (COs) which mimic human brain tissues (Fig. 6a). As we had done in the epithelial and fibroblast cell lines, we first established that neurons in COs of healthy controls exhibited eSGs after a 1-h treatment with two severe energy stresses (Fig. 6b,c). Overall, the eSGs were smaller in size than the conventional SGs induced by arsenite. We found that the severe energy stresses consistently led to a significant decline in ATP concentration in the COs (Fig. 6d). Notably, we found that more glycolytic blockage-induced eSGs were formed in neurons in C9-ALS COs than in neurons of healthy controls (Fig. 6b,c). Since OXPHOS becomes the dominant energy source when glycolysis is blocked, this result is consistent with the reduced OXPHOS activity in C9-ALS neurons and further suggests that such a defect renders patients’ cells more vulnerable to disturbances in glycolytic activity and eSG formation. In line with these observations, we also noted that the ATP concentrations of C9-ALS COs that had been subjected to glycolytic blockage were reduced to a larger extent than were those in healthy controls’ COs (Fig. 6d). However, unlike the situation we observed for glycolytic blockage, OXPHOS inhibition in low-glucose states induced comparable levels of eSGs in healthy and C9-ALS neurons, further confirming that the impaired OXPHOS activity was responsible for the reduced energetic flexibility in the C9-ALS neurons (Fig. 6b,c). Consistently, the ATP concentrations in both healthy and C9-ALS COs were decreased to a similar level when OXPHOS was inhibited (Fig. 6d). Thus, these data suggest that eSG can be induced in CO neurons that have been exposed to severe energy stress and that C9-ALS CO neurons are more prone to eSG formation when glycolysis is inhibited because of their reduced OXPHOS activity. We also observed similar phenotypes in C9-ALS patients’ iPSC-derived motor neurons (MNs). Glycolytic blockage induced eSG formation in control MNs; however, more eSGs were found in C9-ALS MNs under the same conditions (Extended Data Fig. 7a,b). Consistently, the glycolytic blockage reduced the ATP levels in C9-ALS MNs to a larger extent than in the control MNs (Extended Data Fig. 7c).

**Fig. 6.**
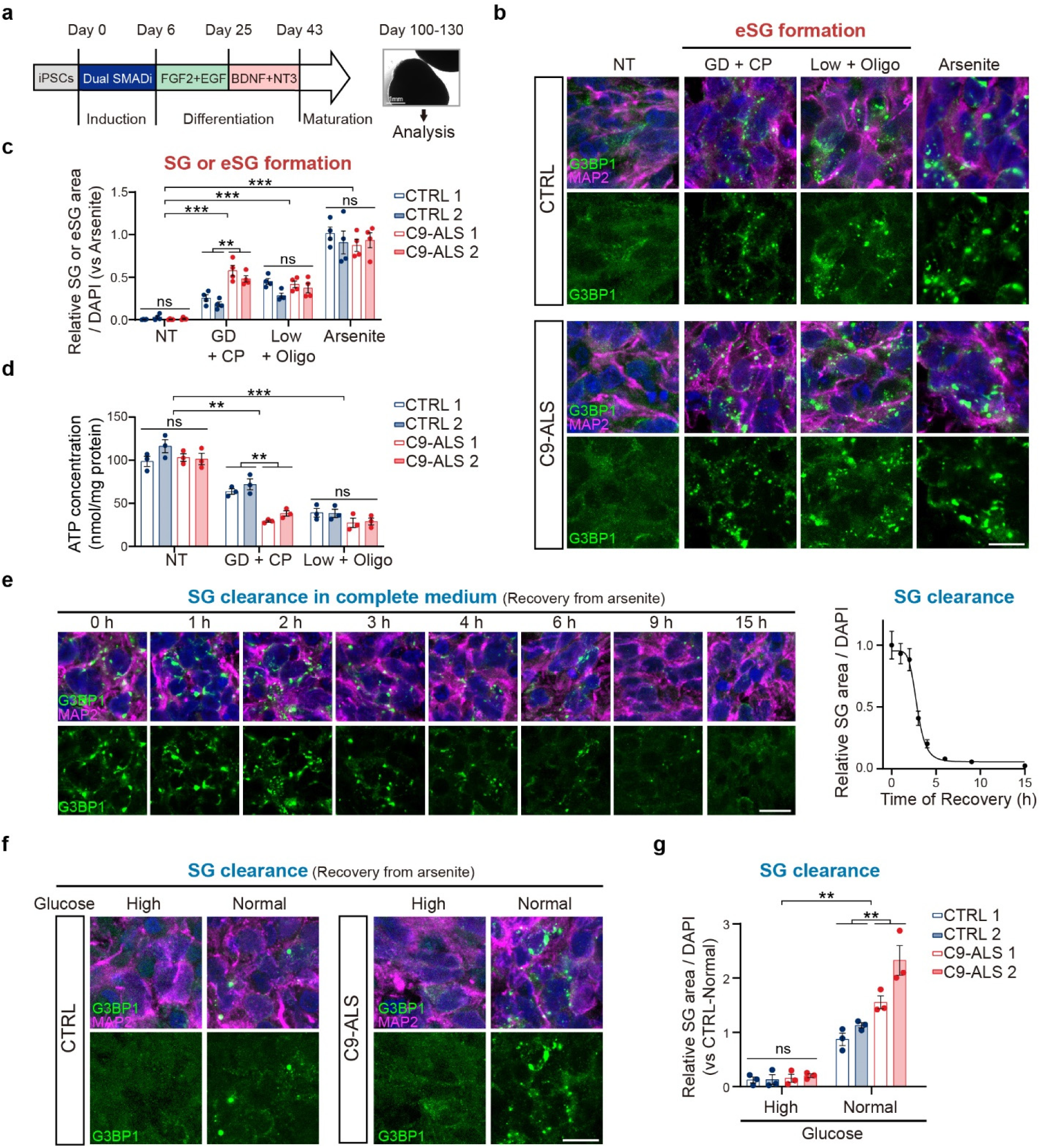
Disrupted eSG formation and SG clearance in C9-ALS patients’ cortical organoids with energy stresses. **a**, A schematic overview of the procedure for generating cortical organoids (COs, 100 to 130 days old) from healthy controls (CTRL) or C9orf72-linked ALS patients’ (C9-ALS) iPSCs. **b**,**c**, Two healthy control COs and two C9-ALS COs treated with either arsenite or the indicated severe energy stress for 1 h were subjected to immunostaining for G3BP1 and a neuronal marker MAP2. Representative images of COs (CTRL 1 and C9-ALS 1) are shown (**b**). The relative granule area per cell (**c**) of SGs or eSGs in COs treated with arsenite or severe energy stresses, respectively, is shown (n = 4). **d**, Intracellular ATP concentrations in healthy controls and C9-ALS COs treated as in (b) (n = 3). **e**, Healthy human COs treated with arsenite for 1 h to trigger SG formation were allowed to recover in complete medium for the indicated time. The persistent SGs in CO neurons were visualized by G3BP1 and MAP2 IF and quantified as the relative SG areas per cell, normalized to the value at time point 0 (n = 3). **f**,**g**, Healthy control and C9-ALS COs after recovery from arsenite treatment in medium with one of the indicated glucose concentrations for 24 h. COs were visualized by G3BP1 and MAP2 IF. Representative images of COs (CTRL1 and C9-ALS1) are shown (**f**). Quantifications of persistent SGs in two healthy controls’ COs or two C9-ALS COs in physiologically high- or normal-glucose medium (**g**) (n = 4). Nuclei were visualized by DAPI staining (blue). Data are shown as means ± SEM, analyzed by unpaired two-sided Student’s *t*-test. * P < 0.05; ** P < 0.01; *** P < 0.001; ns, not significant. Scale bars, 10 μm.

Next, we examined whether the clearance of conventional SGs was also disrupted in C9-ALS patients’ iPSC-derived CO neurons and MNs when moderate energy stresses were present. To this end, we first measured the clearance rate of conventional SGs in healthy human COs and MNs under normal culture conditions. It took 3 h for the MNs to disassemble SGs following arsenite removal, indicating that the clearance of SG in MNs was slightly slower than that in HeLa cells (Extended Data Fig. 7d). In contrast to the 2D cultured cells, however, it took more than 12 h for the CO neurons to disassemble SGs, suggesting that the clearance of SGs in normal human neurons in the 3D tissue is significantly limited (Fig. 6e). We found that the clearance of SGs was significantly delayed in healthy CO neurons even under physiologically normal glucose conditions, as compared to the high glucose group, suggesting that the extracellular glucose concentrations are critical for neurons to disassemble SGs efficiently (Fig. 6f,g). Notably, the SG clearance rates were even slower in C9-ALS patients’ CO neurons under physiologically normal glucose conditions than in healthy control neurons, indicating that the intrinsic defects in energy metabolism reduced the SG clearance capacity in the C9-ALS patients’ CO neurons (Fig. 6f,g). When we examined the SG clearance rates in healthy MNs, similar reductions in SG clearance were observed in healthy neurons cultured under the physiologically normal glucose condition when compared with the high-glucose groups (Extended Data Fig. 7e,f). Moreover, as was observed for the CO neurons, the SG clearance efficiency was further reduced in C9-ALS MNs (Extended Data Fig. 7e,f). These data suggest that an adequate glucose concentration is crucial for effective clearance of conventional SGs in neurons, and ALS/FTD patients’ neurons with defective energy metabolism are more prone to harboring long-lived SGs because of their reduced capacity to clear SGs.

## Discussion

The formation of RNP granules and their dynamics are governed by multifactorial principles in both the assembly and clearance processes. The fate of RNP granules depends on the nature and extent of the energy stresses exerted. Here we demonstrate that energy stress induces the formation of a type of RNP granule distinct from the conventionally studied stress granules and that energy stress plays a critical role in regulating the dynamics of these granules in a dose-dependent manner. We propose a two-tier model for energy-dependent regulation of RNP granule dynamics, in which moderate stresses reduce the ATP-dependent RNA-binding capacity of eIF4A and SG clearance efficiency, whereas severe stresses reduce both eIF4A and mTOR activity, leading to 4EBP1 dephosphorylation and eSG formation (Fig. 7).

**Fig. 7.**
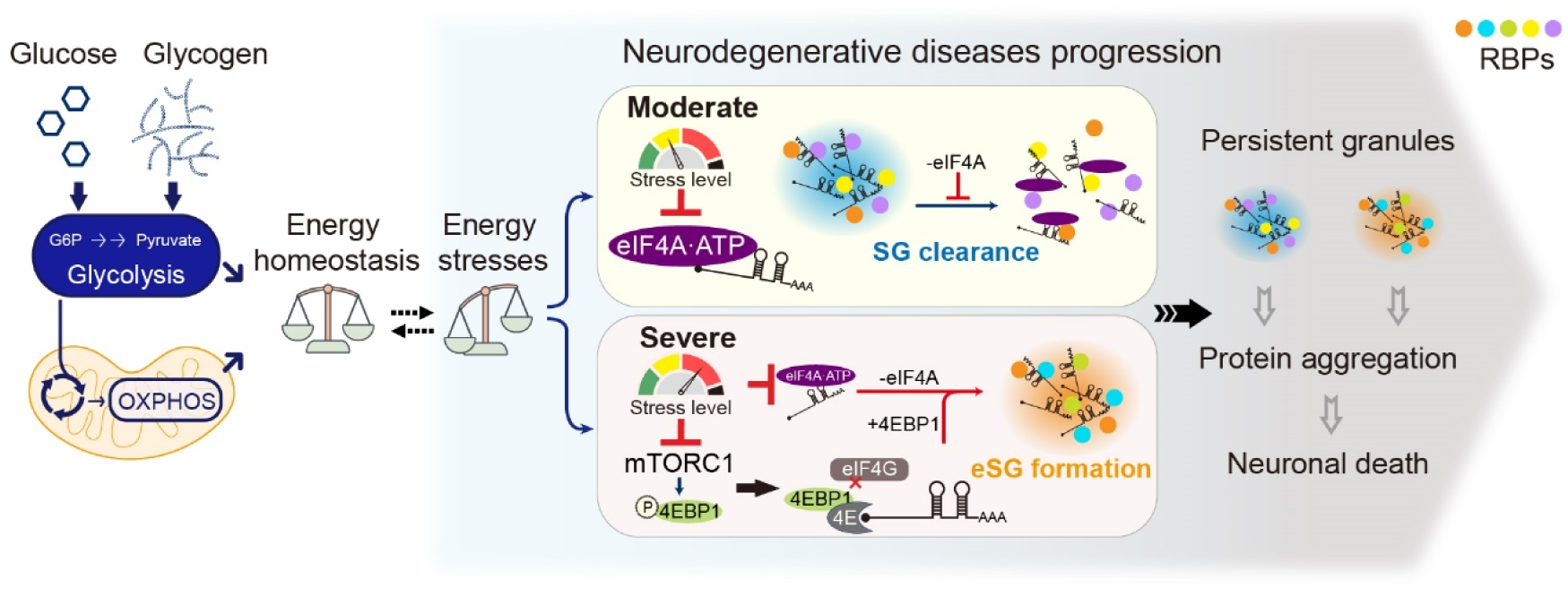
Summary of energy stress-regulated granule dynamics in mammalian cells. Both physiological extracellular glucose and cell-intrinsic glycogen sustain cellular energy production through glycolysis and mitochondrial OXPHOS. During aging or at the early stage of the pathogenesis processes of neurodegenerative diseases, reduced metabolic flexibility causes different levels of energy stresses. On the one hand, moderate stress reduces the ATP-dependent RNA-binding capacity of eIF4A and SG clearance efficiency; on the other hand, severe stress reduces both eIF4A and mTOR activity, leading to 4EBP1 dephosphorylation and eSG formation. The dysregulated SG/eSG dynamics eventually cause the formation of persistent granules, protein aggregation, and neuronal death.

Emerging evidence suggests that the assembly and composition of stress-induced RNP granules are diverse and stress-specific^64–66^. We found that the 4EBP1-dependent eSGs are smaller in size than the arsenite-induced, eIF2α-dependent conventional SGs; furthermore, the shape and assembly dynamics of eSGs also differed from those of SGs. Although eSGs and SGs apparently share protein components that are common building blocks for stress-induced granules, eSGs also contain unique proteins with a nuclear localization and functions. It has been reported that dephosphorylation of 4EBP1 induced by selenite or hydrogen peroxide treatment causes formation of unconventional stress granules lacking eIF3^44, 67, 68^. Also, glucose starvation induces eIF3-lacking granules in yeast^69^. However, we found that the 4EBP1-dependent eSGs that form in mammalian cells under severe energy stress contain eIF3 (Extended Data Fig. 5e,f). Moreover, previous studies have suggested that inhibition of eIF4A can induce stress granules featuring condensed RNAs, which may mediate LLPS independent of G3BP proteins^60, 68^. We have now confirmed that RNA is more condensed in eSGs than in arsenite-induced conventional SGs (Fig. 5). However, such energy stress-induced RNA condensation still requires G3BP proteins to trigger eSG formation. Therefore, the unique induction criteria, protein composition, and underlying mechanistic features of eSGs suggest they are a new subtype of SGs distinct from granules induced by either 4EBP1 activation or eIF4A inhibition.

Standard cell culture media contain excess glucose, which is beneficial for cell maintenance and acts as an energy reservoir to prevent energy disturbances. When we set the glucose concentrations in our culture media within the physiological range, we observed that the extracellular glucose was a major determinant of eSG/SG dynamics: First, physiologically high glucose (5 mM) prevented eSG formation and sustained SG clearance in cells. Second, physiologically normal glucose (1 mM) slightly reduced the intracellular ATP levels but did not cause enough energy stress to disturb the eSG/SG dynamics. However, when another cellular energetic pathway, such as glycogen metabolism or mitochondrial OXPHOS, was also inhibited in the presence of physiologically normal levels of glucose, the cells encountered moderate energy stresses and could no longer clear the SGs. This phenomenon suggests that physiologically normal glucose-fueled glycolysis cannot act as a functional reservoir to serve as a cellular energy supply in response to heightened energy demand under stress, and such reduced metabolic flexibility leads to a failure to clear the SGs. Finally, physiologically low glucose (0.25 mM) alone caused moderate energy stress and delayed SG clearance. In addition, eSGs were formed when glycogenolysis or OXPHOS was blocked under low-glucose conditions, suggesting that a further reduction in metabolic flexibility leads to more persistent eSGs/SGs by enhancing their formation and suppressing the clearance of these granules.

Our studies have provided a mechanistic model for the regulation of eSG formation that implicates both translation and ATP-dependent processes. We found that eSG formation and SG clearance in response to different degrees of energy stress did not involve changes in the phosphorylation of eIF2α, consistent with a previous report that energy stress does not alter the phosphorylation of eIF2α in mammalian cells^25^. Next, we found that the induction of eSGs under severe energy stress, with a 50% reduction in cellular ATP, required the mTOR-4EBP1 axis but not AMPK, consistent with the notion that mTOR can directly sense cellular ATP independent of AMPK^70, 71^. When we reduced cellular ATP, mTOR was inhibited, leading to dephosphorylation and activation of 4EBP1. Activated 4EBP1 then bound eIF4E and blocked the recruitment of eIF4G to the eIF4F complex, an essential step in translation initiation. 4EBP1 was an essential driver for eSG formation, because depletion of this protein dramatically eliminated eSG formation; however, forced activation of 4EBP1 in energy-replete cells did not induce eSGs, suggesting that 4EBP1 activation alone is insufficient to induce eSGs and that additional ATP-sensitive pathways may contribute to eSG formation. Multiple ATP-dependent proteins, including protein chaperones and nucleotide helicases, are closely tied to SG dynamics^28^. Among them, eIF4A is the most abundant DEAD-box RNA helicase in HeLa cells and prevents the formation of stress granules through its ATP-dependent RNA-binding capacity^60^. We found that eIF4A also played a role in preventing eSG formation when this enzyme was supported by an adequate level of intracellular ATP.

Moderate energy stresses, with a 25% reduction in cellular ATP, were sufficient to disrupt the RNA-binding ability of eIF4A, leading to G3BP1 and mRNA condensation into eSGs/SGs. However, such energy stress-induced RNA condensations was not sufficient to promote eSG assembly without G3BP proteins. In fact, our data indicate that both 4EBP1 activation and eIF4A inhibition are required for eSG formation, suggesting that the multivalency required for LLPS and eSG assembly under physiological pathways is regulated by multiple pathways. Notably, in addition to eIF4A, other ATPases have also been shown to regulate SG dynamics. For example, recent studies have suggested a role for VCP, an AAA ATPase functioning as a segregase, in eliminating SGs through either the ubiquitin-proteasome or autophagy pathway^14, 72^. Thus, reductions in ATP levels may regulate eSG/SG dynamics by affecting additional ATPases.

Using both 3D and 2D neuronal culture models, we have found that different degrees of energy stress induce eSG formation or delay SG clearance in human organoids and motor neurons. The energy metabolism in the brain must be fine-tuned to reach a balance between the high energy demands of neurons and the restriction in nutrient supply caused by the existence of the blood-brain barrier. It has been proposed that in neurons, glycolysis is responsible for basal energy demands, whereas OXPHOS is responsible for activity-induced energy demands^73, 74^. Consistently, we observed that both glycolysis and OXPHOS were crucial for the maintenance of energy homeostasis and normal eSG/SG dynamics in neurons. The brain uses both circulating glucose and stored glycogen for glycolysis and OXPHOS^75^. Thus, glycogen is an indispensable source for energy generation and buffering in the brain. Whereas glycogen has long been considered to be stored in astrocytes, recent studies have suggested that neurons also have an active glycogen metabolism that supports neuronal synaptic plasticity and resistance to stress^76–80^. Consistent with this concept, we found that glycogen was critical for supplying energy via glycolysis and for eSG/SG dynamics in neurons in the presence of physiological levels of glucose. Brain glycolysis is usually reduced during normal aging or in NDDs^31, 81^. Moreover, abnormal glycogen deposits in the human brain are observed with aging and in multiple NDDs, including ALS, Alzheimer’s disease, Parkinson’s disease, and Lafora disease^35, 78, 82–84^. Recent studies have indicated that upregulated glycolysis, achieved by either a high-glucose diets or genetic or pharmacological activation of glycolytic enzymes, might mitigate the disease phenotype in animal models or patients with NDDs, including ALS, Parkinson’s disease, and Huntington’s disease^85–87^. Therefore, the disrupted glycolysis and glycogenolysis observed in age-related NDDs may compromise neuronal metabolic flexibility, leading to heightened energy stress in the affected cells. In addition to glycolytic deficits, disrupted mitochondrial OXPHOS is another common pathological feature that contributes to the disrupted energy metabolism in NDDs. We found that OXPHOS is also critical for adequate energy supply under physiologically normal glucose conditions, suggesting that impairment of mitochondrial OXPHOS also contributes to the buildup of energy stress in NDDs.

Persistent SGs are thought to be associated with the proteinaceous pathology and neuronal cell death seen in multiple NDDs, including ALS and FTD. Disruptions in glycolysis and mitochondrial function in ALS/FTD occur in presymptomatic individuals and precede disease onsets^34, 88^. Our data support a pathogenic model of ALS/FTD in which disrupted energy metabolism and subsequent energy stresses lead to neuronal death (Fig. 7). In principle, aging-related metabolic declines cause a reduction in energetic flexibility, an early event in the disease that exerts energy stress on the affected neurons. Specifically, neurons exposed to either severe or moderate energy stress are more likely to harbor persistent RNP granules resulting from either eSG formation or reduced SG clearance. Furthermore, perturbation of RNP granule dynamics can precipitate the gain- or loss-of-function toxicity of disease-related proteins, such as TDP-43 and FUS, eventually leading to protein aggregations and neuronal death. Thus, our results have established a mechanistic connection among energy stress, metabolic flexibility, and granule dynamics that could have broad implications for understanding the energetic basis of neurodegenerative diseases.

## Methods

### Plasmids and oligonucleotides

The mCherry-G3BP1 plasmid was constructed by PCR amplification to generate a complementary DNA (cDNA) encoding mCherry-G3BP1 and then ligated into pEGFP-C1 vector digested with AgeI/BamHI to remove EGFP. The AU1-tagged wild-type and two mutant (S2215Y and R2505P) mTOR constructs were gifts from Fuyuhiko Tamanoi (Addgene) ^59^. The V5-tagged human GLUT1 construct was a gift from Wolf Frommer (Addgene) ^89^. cDNA of N-terminal Flag-tagged wild-type human eIF4A1 was amplified by PCR and subcloned into the pLenti-CMV-Puro-DEST (w118-1) vector ^90^ using the Gateway cloning system (Thermo Fisher). The TIMMDC1, NDUFS5, and 4EBP1 shRNA constructs were purchased from Sigma: human TIMMDC1 (TRCN0000154913); human NDUFS5 (TRCN0000036643); human 4EBP1 #1 (TRCN0000040203); human 4EBP1 #2 (TRCN0000040206). The Cy5-oligo-dT_20_ probe was purchased from IDT. The gRNAs targeting human AMPK α1 or α2 described previously were cloned into the gRNA/Cas9 expression vector pLenti-CRISPR v2 (Addgene) ^91, 92^.

### Cell lines

HeLa, MEF, U2OS, and HEK293 cells were maintained in Dulbecco’s Modified Eagle’s Medium (DMEM) supplemented with 10% FBS and 1% penicillin-streptomycin (10,000 units/mL penicillin and 10 mg/mL streptomycin). The immortalized wild-type MEFs were generated from wild-type C57BL/6 mice ^93^. The eIF2α S51A MEFs were previously described ^94^. RPE1 cells were maintained in DMEM/F12 supplemented with 10%FBS, 0.01 mg/mL hygromycin B, and 1% penicillin-streptomycin. G3BP1/G3PB2 DKO U2OS cells were described previously ^4, 95^. All cell lines were cultured in a humidified cell incubator at 37°C in a 5% CO_2_ atmosphere.

The iPSCs were obtained from the National Institutes of Health Cell Repositories and maintained on Matrigel (BD)-coated 6-well plates in mTeSR Plus medium (STEMCELL Technologies) using a feeder-free culture protocol. Cells were cultured at 37°C in a hypoxic (5% O_2_) incubator.

### Cell transfection and treatment

HeLa cells stably expressing mCherry-G3BP1 were established by transfection of the mCherry-G3BP1 construct using Lipofectamine 3000 (Thermo Fisher) according to the manufacturer’s instructions and subsequent multiple rounds of G418 (Thermo Fisher) selection. A single colony was picked and expanded. Cells were then maintained in the standard complete medium without G418 for at least ten passages before further analysis. For the 4EBP1 knockdown in HeLa cells, the plasmid encoding two independent shRNAs targeting 4EBP1 or a non-targeting control was transfected into HeLa cells using Lipofectamine 3000. The transfected cells were analyzed 72 to 96 h after transfection. HeLa cells with stably knocked down NDUFS5 or TIMMDC1 were constructed by lentiviral infection. The CRISPR-Cas9 system was used to generate the AMPK α1/α2 double-knockout HeLa cells through lentiviral infection. For lentivirus preparation, HEK293 cells were co-transfected with a lentiviral vector plus two packaging plasmids, pSPAX2 and pMD2.G, using Lipofectamine 3000. Forty-eight hours after transfection, lentiviral supernatant was collected and subsequently filtered through a 0.45-μm cellulose acetate filter. For all the lentivirus infections, HeLa cells were plated at 3 × 10^4^ cells per well in 6-well plates and then supplemented with viral medium for 72 h. Subsequently, cells were treated with puromycin (Thermo Fisher) for 72 h at a final concentration of 1.5 μg/ml. After puromycin selection, single colonies were picked, and cells were maintained in the standard complete medium without puromycin for at least ten passages to minimize the potential effects of puromycin on stress granule dynamics before further analysis. The resulting cell lines were verified by either sequencing the targeted locus or probing the targeted protein through immunoblot analyses.

For glycolysis inhibition, HeLa, MEF, U2OS, or RPE1 cells were first rapidly rinsed with PBS and cultured in GD medium [glucose-free DMEM (Thermo Fisher) containing 10% dialyzed FBS (Thermo Fisher)] supplemented with either 50 μM CP91149 (Selleckchem) (GD + CP) or 25 mM 2DG (Sigma) (GD + 2DG), or in the standard culture medium supplemented with 25 μM shikonin (Sigma) (SKN) for 1 h. For PPP inhibition, HeLa cells were rinsed with PBS and cultured in the normal culture medium supplemented with 100 μM 6-aminonicotinamide (Sigma) (6AN) for 1 h. For the assays using the physiological range of glucose, HeLa cells were rinsed with PBS and cultured in the GD medium supplemented with glucose at indicated concentrations for 1 h, with or without 50 μM CP91149 or 2 μM oligomycin (Cell Signaling Technology). For the glutamine deprivation, cells were rinsed with PBS and cultured in phenol-free DMEM (without glucose or glutamine) (Thermo Fisher) supplemented with 10% dialyzed FBS and glucose at indicated concentrations for 1 h. Conventional SGs were induced in cells treated with 250 μM arsenite (Sigma) for 1 h. For the SG clearance analysis, HeLa cells harboring arsenite-induced conventional SGs were briefly washed with PBS and cultured in complete or various starvation media to recover for 1 h.

For the induction of eSGs in MNs or COs, neurons or organoids were rapidly washed with PBS, then maintained in the neuronal GD medium [glucose-free neurobasal medium (Thermo Fisher) supplemented with 1X Glutamax (Thermo Fisher)], in combination with either 50 μM CP91149 (GD + CP) or 0.25 mM glucose (Sigma) plus 2 μM oligomycin (Low + Oligo), for 1 h before harvesting. Conventional SGs in MNs or COs were induced by 1 h administration of 250 μM arsenite in the normal culture medium. For SG clearance analysis in MNs or COs, neurons or organoids harboring arsenite-induced conventional SGs were briefly washed with PBS and allowed to recover in the normal culture medium or the neuronal GD medium supplemented with either 5 mM (High) or 1 mM (Low) glucose.

Liposomal ATP (ATPsome) was purchased from Encapsula NanoScience. Lyophilized proliposomes comprising a 7:3 molar ratio of phosphatidylcholine (PC): phosphatidylserine (PS) was dissolved in water immediately before use to form 100 nm liposomes containing 1 mM ATP. To rescue the energy deficiency in cultured cells under the treatments described above, ATPsome (10 μM ATP) was added to the medium twice during the 1 h treatment: once at the beginning of treatment and a second time at 30 min. For the Dm-αKG rescue experiments, cells were treated with the indicated media in the presence of 4 mM Dm-αKG.

### Cortical organoid and motor neuron generation

The method used for converting the human iPSCs into motor neurons was based on a previous report with minor modifications ^96^. Briefly, on day 0, iPSCs were suspended and transferred into one Matrigel-coated well of a 6-well plate with 10 μM Rock inhibitor Y27632 (Stemgent). On day 1, the medium was changed to neuronal medium [a mixture of 50% Neurobasal medium (Thermo Fisher) and 50% DMEM/F12 medium (Thermo Fisher) with 1X Glutamax (Thermo Fisher), 0.5X N2 and B27 supplement (Thermo Fisher), and 0.1 mM ascorbic acid (Sigma)], supplemented with 3 μM CHIR99021 (Stemgent), 2 μM SB431542 (Stemgent), and 2 μM DMH-1 (Tocris) for 6 d. Cells were then dissociated using Dispase (STEMCELL Technologies) and seeded into Matrigel-coated 10-cm dishes in neuronal medium plus 1 μM CHIR99021, 2 μM DMH-1, 2 μM SB431542, 0.1 μM retinoic acid (RA) (Sigma), and 0.5 μM purmorphamine (Stemgent). On day 13, cells were digested with Dispase and placed in 10 cm ultra-low adhesion plates (Corning) in neural medium plus 0.5 μM RA and 0.2 μM purmorphamine. On day 21, cells were dissociated with 1X Accutase (Thermo Fisher) and seeded at a density of 2 × 10^6^ cells per well (6-well plates) or 0.5 × 10^6^ cells per well (24-well plates) into PDL/Laminin (Sigma)-coated plates with neuronal medium plus 0.5 μM RA, 0.2 μM purmorphamine, and 0.1 μM Compound E (Sigma). Cells were cultured for additional 12 to 14 days with medium change every 2 days before experimental analysis.

Cortical organoids were generated as previously described with minor modifications ^97, 98^. Briefly, the iPSCs were subcultured in the vitronectin (Thermo Fisher)-coated 10 cm dishes with Essential 8 medium (Thermo Fisher). After reaching ∼70% confluency, cells were treated with 1% DMSO for 24 h. Then, on day 0, cells were washed with PBS and incubated with 4 mL Accutase for 7 min at 37 °C in a 5% CO_2_ incubator and dissociated into single cells with gentle pipetting. Cells were collected with centrifugation in a 50-mL conical tube and washed with Essential 8 medium. Cells were suspended in Essential 8 medium containing 10 μM ROCK inhibitor Y27632, and cell numbers were determined using a TC20 cell counter (Bio-Rad). AggreWell 800 (STEMCELL Technologies) plates containing 300 microwells were used to generate uniformly sized spheroids. 3 × 10^6^ cells were added into each AggreWell 800 well and subjected to centrifugation at 100 g for 3 min to distribute the cells at 10,000 cells per microwell. After being cultured at 37 °C in a 5% CO_2_ incubator for 24 h, the iPSCs-derived spheroids were resuspended by pipetting using a 7-mL plastic transfer pipette. The spheroid suspensions were placed in a 40-μm cell strainer (Fisher Scientific), and the retained spheroids were collected by washing the strainer with Essential 6 medium (Thermo Fisher). Harvested spheroids then were transferred into 10-cm ultra-low adhesion plates in Essential 6 medium containing dual SMAD pathway inhibitors [100 nM LDN193189 (STEMCELL Technologies) and 10 μM SB431542] and 2.5 μM XAV939 (Cayman). From day 2 to day 6, the medium was changed daily. On day 6, the medium was replaced with CO neuronal medium consisting of Neurobasal A (Thermo Fisher) plus 1X B27 supplement minus vitamin A (Thermo Fisher) and 1X Glutamax in the presence of 20 ng/mL EGF (PeproTech) plus 20 ng/mL FGF2 (PeproTech). CO neuronal medium supplemented with EGF/FGF2 was changed daily for 10 days and then every two days for additional 9 days. On day 25, the medium was replaced with CO neuronal medium supplemented with 20 ng/mL BDNF (PeproTech) and 20 ng/mL NT3 (PeproTech); the medium was changed every two days. From day 43 on, the medium was replaced every 4 days with CO neuronal medium without growth factors. The plates were placed on an orbital shaker rotation at 40 rpm to minimize organoid fusion.

### Cryosection of cortical organoids

COs were quickly rinsed with PBS and fixed for 24 h at 4 °C in 4% paraformaldehyde. COs were then washed three times with PBS and transferred into 30% sucrose for 72 h. Organoids were embedded using a 1:1 mix of OCT compound (Tissue Tek) with 30% sucrose and snap-frozen on dry ice and stored at -80 °C. Organoids were cryosectioned at 16-μm thickness using a CryoStar NX70 cryostat (Thermo Fisher) and collected on Superfrost Plus glass slides (Fisher Scientific). Sections were stored at -80 °C for subsequent analysis.

### Cell viability assay

Cell viability in HeLa cells treated with pro-death stressors was measured by staining cells with annexin V AlexaFluor-488 conjugate (Thermo Fisher) as described previously ^39^. In brief, 2 × 10^5^ cells from each group were incubated with 200 µl of stain for 30 min at room temperature in the dark, followed by fluorescent microscopy examination (Nikon Eclipse TS100). Annexin V-positive cells were counted in three randomly selected fields of view using a 20X objective. For each group, at least 500 cells from three independent experiments were analyzed.

### Immunofluorescence (IF) and fluorescence in situ hybridization (FISH)

For immunofluorescence analysis, cells grown on coverslips were rapidly rinsed with PBS and then fixed in 4% paraformaldehyde in PBS for 20 min at room temperature. Fixed cells were then permeabilized with 0.1% Triton X-100 in PBS and treated with blocking buffer (1X PBS, 3% BSA, and 0.1% Tween-20) for 30 min at room temperature. Cells were incubated with primary antibodies in blocking buffer overnight at 4°C and washed with 1X PBS supplemented with 0.1% Tween-20. Then cells were incubated with fluorescently conjugated secondary antibodies in blocking buffer for 1.5 h at room temperature and washed with 1X PBS supplemented with 0.1% Tween-20. Coverslips were mounted with ProLong Diamond with DAPI (Thermo Fisher).

For immunofluorescence analysis of organoid cryosections, slides were washed with PBS for 5 min to remove the excess embedding medium and then treated with CO blocking buffer (1X PBS, 4% BSA, and 0.3% Triton X-100) for 1 h at room temperature. Sections were incubated with primary antibodies in CO blocking buffer overnight at 4°C. Primary antibodies were washed off with PBS supplemented with 0.3% Triton X-100. Sections were then incubated with fluorescently conjugated secondary antibodies in CO blocking buffer. After incubation for 1.5 h at room temperature, slides were washed with 1X PBS supplemented with 0.3% Triton X-100 and mounted with ProLong Diamond with DAPI.

The FISH assay was performed as previously described with minor modifications ^99^. Briefly, cells grown on coverslips were fixed in 4% paraformaldehyde in PBS for 10 min at room temperature, followed by permeabilization in ice-cold 70% ethanol overnight at 4°C. Cells were washed twice with wash buffer (2X SSC and 10% deionized formamide) and prehybridized at 37°C for 10 min in hybridization buffer (10% deionized formamide, 2X SSC, 50 μg/mL BSA, 100 mg/mL dextran sulfate, 1 mg/mL yeast tRNA, and 2 mM vanadyl sulfate ribonucleosides). Cells were then incubated with the hybridization buffer containing 100 nM Cy5-oligo-dT_20_ probes at 37°C overnight, followed by washing three times using pre-warmed wash buffer for 10 min.

Primary antibodies used for the immunofluorescent staining were as follows: anti-HUR (Santa Cruz Biotechnology, sc-5261), anti-eIF4G (Cell Signaling Technology, 2469), and anti-TDP-43 (Proteintech Group, 10782-2-AP), anti-G3BP1 (BD Bioscience, 611126), anti-V5 tag (Thermo Fisher, 460705), anti-TIA1 (Proteintech Group, 12133-2-AP), anti-PABP (Santa Cruz Biotechnology, sc-32318), anti-EIF2A (Proteintech Group, 11233-1-AP), anti-ACP1 (Proteintech Group, 22214-1-AP), anti-TNPO3 (Thermo Fisher, MA5-34790), anti-NUP107 (Proteintech Group, 19217-1-AP), anti-NBN (Proteintech Group, 55025-1-AP), anti-hnRNPM (Proteintech Group, 26897-1-AP), anti-MRE11 (Cell Signaling Technology, 4895), anti-RMI1 (Proteintech Group, 14630-1-AP), anti-4EBP1 (Cell Signaling Technology, 9644), anti-eIF3B (Santa Cruz Biotechnology, sc137214), anti-ChAT (Sigma, AB143), and anti-MAP2 (Abcam, ab32454).

### Live-cell imaging

Live-cell imaging of SGs or eSGs was performed with HeLa cells stably expressing mCherry-G3BP1 using SP8 Confocal Microscope (Leica) equipped with a 63X oil objective. Cells were plated in 35-mm coverslip-bottom dishes (MatTek). Before imaging, cells were rapidly rinsed with PBS, and the medium was replaced with phenol-free DMEM (without glucose or glutamine) (Thermo Fisher) containing 10% dialyzed FBS, 4 mM glutamine, and 50 μM CP91149 (GD + CP) or the phenol-free DMEM supplemented with 10% FBS, 25 mM glucose, 4 mM glutamine, and 250 μM arsenite (Arsenite). Cells were imaged every 2 min for 1 h at 37°C, 5% CO_2_ in an environmental control chamber.

### Stress granule isolation

SGs or eSGs were purified as described previously ^51^. In short, HeLa cells stably expressing mCherry-G3BP1 were grown to 90% confluency in five 15-cm culture dishes for each group. One hour before stress treatment, the cell culture medium was replaced with the fresh complete medium. Cells were then treated with stresses as described above. After the stresses, cells were directly scraped down without PBS wash and pelleted by centrifugation at room temperature for 3 min. After aspirating the remaining medium, the pellets were snap-frozen in liquid nitrogen and subjected to the freeze and thaw cycle three times. Cells were resuspended in the lysis buffer [50 mM Tris HCl, pH 7.4, 100 mM potassium acetate, 2 mM magnesium acetate, 0.5 mM DTT, 50 µg/mL heparin, 0.5% NP40, 0.5 U/µl RNasIN (Promega) RNase inhibitor, and 1X protease inhibitor cocktail (Sigma)] and passed through a 25G needle for five times, followed by a brief sonication at the low power (5 s pulse with 15 s recovery) on ice for a total of 2 min. The lysates were cleared by centrifugation at 1,000 g for 5 min. The supernatants were transferred into a clean tube and subjected to centrifugation at 18,000 g for 15 min. The pellets containing SG cores were resuspended in the lysis buffer by firmly pipetting and then pre-cleared by incubating with protein A/G Dynabeads (Thermo Fisher) at 4°C for 30 min. After removing Dynabeads, lysates were incubated with 2.5 µg anti-mCherry antibody (Thermo Fisher, PA5-34974) and rotated overnight at 4°C. After spinning at 18,000 g for 15 min at 4°C, the supernatant was removed to discard the unbounded antibody. Pellets were resuspended and incubated with protein A/G Dynabeads at 4°C for 3 h. The Dynabeads were washed three times with lysis buffer, once with washing buffer 1 (lysis buffer containing 2 M Urea), once with washing buffer 2 (lysis buffer containing 300 mM potassium acetate), and three times with 50 mM Tris HCl (pH 7.4). The enrichments of mCherry-G3BP1-positive granules at each step were checked by microscopy.

### Proteomic analysis

The bead-bound proteins were lysed by sonicating in 8 M urea/50 mM triethylammonium bicarbonate (TEAB). They were reduced and alkylated with 10 mM tris (2-Carboxyethyl) phosphine hydrochloride and 40 mM chloroacetamide for 1 hour at room temperature. Protein digestion was carried out with LysC (Fujifilm Wako) in 10 ng/µL (enzyme-to-protein, w/v) at 37°C for 3 h. The proteins were further digested with trypsin (Promega) in 10 ng/µL (enzyme-to-protein, w/v) at 37°C overnight after diluting the urea concentration to 2 M by adding 50 mM TEAB. The digested peptides were desalted with C18 StageTips. The eluted peptides were reconstituted in 0.5% formic acid (FA) after vacuum-drying, and then they were injected into the mass spectrometer. All peptide samples were analyzed on an Orbitrap Fusion Lumos Tribrid Mass Spectrometer with an Ultimate3000 RSLCnano system (Thermo Scientific). They were loaded C18 particles at 8 μL/minute flow rate. They were separated at 0.3 μL/minute flow rate under an increasing gradient of solvent B (0.1% FA in 95% acetonitrile) onto an EASY-Spray analytical column (75 μm × 50 cm, Thermo Scientific) packed with 2-μm C18 particles, with 2.0 kV voltage operation (total run time: 120 minutes). Mass spectrometry analysis was completed in data-dependent acquisition with a mass-to-charge ratio (m/z) range of 300–1,800. MS1 (precursor ions) was acquired with 120,000 resolution at 200 m/z. MS2 (fragment ions) was acquired with 30,000 resolution at 200 m/z using a higher-energy collisional dissociation with 32% collision energy. Automatic gain controls for MS1 and MS2 were set to 1,000,000 and 50,000 ions, respectively. Maximum ion injection times for MS1 and MS2 were set to 50 and 100 milliseconds, respectively. Isolation window was set to 1.6 m/z with a 0.4 m/z offset. Dynamic exclusion was set to 30 seconds with a 7-ppm mass window. Internal calibration was conducted with the lock mass (m/z 445.1200025). Proteome Discoverer (v2.4.1.15, Thermo Scientific) was used for quantitation and identification of proteins. The tandem mass spectrometry data were then searched using SEQUEST algorithms against a human UniProt database (released in Jan. 2021). The search parameters and methods used were as follows: a) trypsin including 2 maximum missed cleavage; b) precursor mass error tolerance of 10 ppm; c) fragment mass error tolerance of 0.02 Da; d) carbamidomethylation (+57.02146 Da) at cysteine as fixed modifications; and e) oxidation at methionine (+15.99492 Da), methionine loss (−131.040485 Da) at protein N-terminus, and methionine loss with acetylation (−89.02992 Da) at protein N-terminus as variable modifications. Peptides and proteins were filtered at a 1% false discovery rate (FDR) at the peptide-spectrum match level using the percolator node and at the protein level using the protein FDR validator node, respectively. The protein quantification was performed using the area under the curve of precursors.

### Gene ontology analysis

The lists of the granule proteins acquired from the proteomic analysis were subjected to the functional annotation clustering analysis by using the DAVID (Database for Annotation, Visualization and Integrated Discovery) online resource (https://david.ncifcrf.gov/) ^100^.

### Ribopuromycinylation assay

Ribopuromycinylation assay was performed as described ^60^. HeLa cells were first treated with indicated stresses for 1 h, as stated above. Then, 5 min before harvesting the cells, puromycin was added to the medium at a final concentration of 5 µg/mL. Cells were washed with ice-cold PBS and lysed in RIPA buffer (50 mM Tris pH 8.0, 150 mM NaCl, 0.1% SDS, 1% NP40, and 1% sodium deoxycholate) supplemented with 1X protease inhibitor cocktail, followed by SDS-PAGE and western blotting analysis.

### UV-RNA immunoprecipitation (UV-RIP)

UV-RIP was performed as described with modifications ^99, 101^. HeLa cells grown on 10-cm dishes were subjected to stresses as stated above. Cells were washed with ice-cold PBS, and then dishes were placed in a UV-light box (UV Stratalinker) without the lid followed by irradiation with 250 mJ/cm^2^ at 254 nm UV light. Cells were collected and resuspended in 0.5 mL of cytoplasmic lysis buffer (25 mM HEPES, pH 7.9, 5 mM KCl, 0.5 mM MgCl_2_, and 0.5% NP40) supplemented with 1X protease inhibitor cocktail and 1.6 U/µL RiboLock RNase inhibitor (Thermo Fisher). Lysates were kept on ice for 30 min and centrifuged at 2,500 g for 5 min at 4°C. The supernatants were set aside on ice, and the pellets were further resuspended in 0.5 mL of nuclear lysis buffer (25 mM HEPES, pH 7.9, 10% sucrose, 350 mM NaCl, and 0.01% NP40) supplemented with protease and RNase inhibitors, followed by a vortex for 30 s and 30 min incubation at 4°C with end-over-end mixing. The nuclear and cytoplasmic fractions were then combined and cleared by centrifugation at 20,000 g for 10 min at 4°C. The supernatant was transferred to a clean tube and pre-cleared with 10 µL protein A/G Dynabeads at 4°C for 1 h. For 1 mL of the precleared cell lysate, a 50 μL aliquot was saved for RNA input and another 50 μL for protein input. The remaining supernatants were incubated with either an anti-eIF4A1 polyclonal antibody (Thermo Fisher, PA5-34726) or a rabbit IgG control (Cell Signaling Technology, 2729S) plus protein A/G Dynabeads 4°C for 2 h with gentle inversion. Beads were washed twice with cytoplasmic lysis buffer, twice with nuclear lysis buffer, and twice with a 1:1 mixture of cytoplasmic and nuclear lysis buffers. One-tenth of the washed beads were saved and boiled in 1X SDS sample buffer (50 mM Tris HCl, pH 6.8, 150 mM NaCl, 1% NP40, 1% sodium deoxycholate, 2% SDS, 1% 2-mercaptoethanol, and 12.5 mM EDTA) for protein examination. The remaining beads were incubated with elution buffer (10 mM Tris HCl, pH 8.0, 1 mM EDTA, 1% SDS, and RNase inhibitor) containing 100 µg proteinase K (Thermo Fisher) at 50°C for 30 min with agitation.

### qPCR

RNA extraction, cDNA generation, and qPCR were performed as described previously ^102^. Briefly, total RNAs in the RNA inputs or the UV-RIP elutions were extracted using the TRIzol reagent (Thermo Fisher). 1 µg RNA was reverse transcribed using QuantiTect Reverse Transcription Kit (Qiagen) according to the manufacture’s protocol, followed by PCR amplification with iQ SYBR Green PCR mix (Bio-Rad). Gene expression data were calculated by the ΔΔCt method.

### ATP measurements

ATP was measured using a luciferase-based luminescence assay (Thermo Fisher). HeLa cells or iPSC-derived motor neurons grown on 6-well plates were rapidly rinsed with ice-cold PBS and subjected to ATP extraction with 0.5% TCA (Sigma). After incubation for 15 min at room temperature, the ATP-containing supernatants were transferred to a fresh tube and neutralized by 0.1 M Tris-HCl buffer (pH 7.75). The remaining proteins on the plates were dissolved in RIPA buffer, and the protein concentrations were measured by BCA assay (Thermo Fisher). The ATP in the COs was extracted using the neutral phenol solution as described previously ^103^. Briefly, each CO was transferred into a clean tube and lysed in neutral phenol (Sigma) using a motor homogenizer; then 0.2X volume of chloroform (Sigma) and 0.15X volume of water was added. Tubes were vortexed for 30 s and centrifuged at 10,000 g for 5 min at 4°C. The upper aqueous phases were collected and diluted with water, followed by ATP concentration determination. Proteins in the interphase were precipitated with 3X volume of isopropanol and dissolved in RIPA buffer for BCA assay. The ATP concentration was determined using a standard curve for the luciferase assay and then normalized to the protein concentration.

### Glucose uptake assay

The glucose transport capacities in the GLUT1-expressing cells were determined using a luciferase-based luminescence assay (Promega). Briefly, HeLa cells were transfected with GFP or GLUT1-V5 constructs and allowed to grow for 48 h before analysis. Cells were washed with PBS three times to remove glucose from the culture and then incubated with PBS containing 1 mM 2DG for 5 min. The glucose uptake rates were assessed by the levels of 2DG6P, which was converted from 2DG once it was imported into the cells in the same manner as glucose.

### Mitochondrial respiration assay

Cells were seeded onto a Seahorse XF96 Cell Culture Microplate (Agilent) at a density of 1.0 × 10^4^ cells per well and allowed to adhere overnight at 37°C and 5% CO_2_. Subsequently, the cells were washed with PBS and incubated in the XF base medium (Agilent) supplemented with 4 mM L-glutamine (Thermo Fisher), 1 mM sodium pyruvate (Thermo Fisher), and 25 mM glucose. Mitochondrial OCRs were determined using the Seahorse XF96 Analyzer (Agilent). The following drugs were injected at the final concentrations as indicated: oligomycin (2 µM), FCCP, (1 µM), and rotenone (0.75 µM) and antimycin A (0.75 µM). Total protein was isolated from each well and quantified by BCA assay for normalization.

### Immunoblotting

Cells were collected and lysed in RIPA buffer plus 1X protease inhibitor cocktail, 1 mM PMSF, 1 mM Na_3_VO_4_, 100 mM NaF, and 1X phosphatase inhibitor cocktail (Sigma) on ice. After centrifugation at 22,000 g for 20 min at 4°C, supernatants were saved, and protein concentrations were determined by the BCA assay. Samples were mixed with 5X SDS sample buffer (250 mM Tris, pH 6.8, 750 mM NaCl, 5% NP40, 5% sodium deoxycholate, 10% SDS, 5% 2-mercaptoethanol, and 60 mM EDTA) and boiled for 10 min before SDS-PAGE analysis. After electrophoresis, proteins were transferred to nitrocellulose membranes and then probed with the indicated antibodies. Actin or Tubulin was used as a loading control.

Primary antibodies for immunoblotting were as follows: anti-puromycin (Sigma, MABE343); anti-GLUT1 (Santa Cruz Biotechnology, sc-377228); anti-pAMPKα (Thr172) (Cell Signaling Technology, 50081); anti-AMPKα (Cell Signaling Technology, 5831); anti-pRaptor (Ser792) (Cell Signaling Technology, 2083); anti-Raptor (Cell Signaling Technology, 2280); anti-pACC (Ser79) (Cell Signaling Technology, 3661); anti-ACC (Cell Signaling Technology, 3676); anti-peIF2α (Ser51) (Cell Signaling Technology, 9721); anti-eIF2α (Cell Signaling Technology, 5324); anti-eIF4G (Cell Signaling Technology, 2469); anti-eIF4E (Cell Signaling Technology, 2067); anti-p4EBP1 (Ser65) (Cell Signaling Technology, 9451); anti-4EBP1 (Cell Signaling Technology, 9644); anti-pS6K (Thr389) (Cell Signaling Technology, 9234); anti-S6K (Cell Signaling Technology, 9202); anti-β-tubulin (Cell Signaling Technology, 2128); and anti-β-actin (Santa Cruz Biotechnology, sc-47778).

### m^7^GTP-sepharose pulldown assay

Capped mRNAs were pulled down by m^7^GTP-sepharose (Jena Bioscience) as described with modifications ^44^. Briefly, cells grown on 10-cm dishes were collected in ice-cold PBS and lysed with pulldown lysis buffer (50 mM Tris HCl, pH 7.5, 100 mM NaCl, 1mM EDTA, and 0.5% NP40) plus 1X protease inhibitor cocktail on ice for 30 min. After centrifugation at 18,000 g for 20 min at 4°C, supernatants were transferred into clean tubes, and protein concentrations were determined by the BCA assay. For each group, an aliquot was saved as an input control. The supernatant containing 1 mg of total protein was incubated with prewashed 15 µL suspension of m^7^GTP-sepharose for 2 h at 4°C with rotation. The beads were washed with the pulldown lysis buffer five times, followed by incubation with 1X SDS sample buffer, and boiled for 5 min to elute the cap-bounded proteins.

### Statistical analyses

All the data are presented as means ± SEM. The band intensity in immunoblots was determined using Bio-Rad Quantity One software. For single comparisons, the statistical significance was analyzed using a Student’s *t*-test. For multiple means of comparison, the one-way analysis of variance (ANOVA) was performed, and the statistical significance was analyzed by the Bonferroni post-hoc test. *P* < 0.05 was considered to be statistically significant. The sample size (n) represents biological replicates per experiment. Data were plotted with GraphPad Prism.

Images were captured by an SP8 Confocal Microscope (Leica) equipped with a 63X oil objective using the same settings to allow the comparison of signal intensities between each group. Images were analyzed using Fiji software. For analyses using the mCherry-G3BP1 expressing cells, six fields of view per group were analyzed, and only cells with equivalent expression levels of mCherry-G3BP1 were used for analysis. For comparisons of the percentage of SG positive cells, cells harboring at least five visible SGs/eSGs with diameters > 0.5 µm or one SG/eSG with a diameter > 2 µm were considered as SG/eSG positive cells for each group. For comparisons of the SG/eSG areas in cell cultures, each cell was first segmented, and the average intensity was determined. The granules were identified using the ROI manager tool according to size and contrast thresholding. The total size of granules in each cell was measured. For assessing the condensation of mRNA and G3BP1, cells were segmented individually, and the granules were identified as stated above. The mean intensities of G3BP1 and oligo-dT probes inside each granule ROI were measured. The granules in COs were assessed by dividing the total granule area in each field of view by the total nuclear area based DAPI staining.

## Acknowledgments

This work was supported by grants from NIH (NS074324, NS089616, NS110098, S10OD021844), Packard Center for ALS Research at Johns Hopkins, Muscular Dystrophy Association, and the U.S. Department of Defense. We thank J. Paul Taylor for providing the G3BP1/G3BP2 DKO U2OS cells, NINDS and NIGMS Cell Repository, Johns Hopkins Brain Bank, and VA Biorepository Brain Bank for providing patient cells and tissues, and members of Wang lab for discussion.

## Author contributions

T.W. and X.T. performed and analyzed experiments. Y.J. and C.H.N. performed the proteomic analysis. P.H. helped with cell culturing and data collection. T.W. and J.W. designed the studies and wrote the paper.

## Competing interests

The authors declare no competing interests.

---

**Extended Data Fig. 1.**
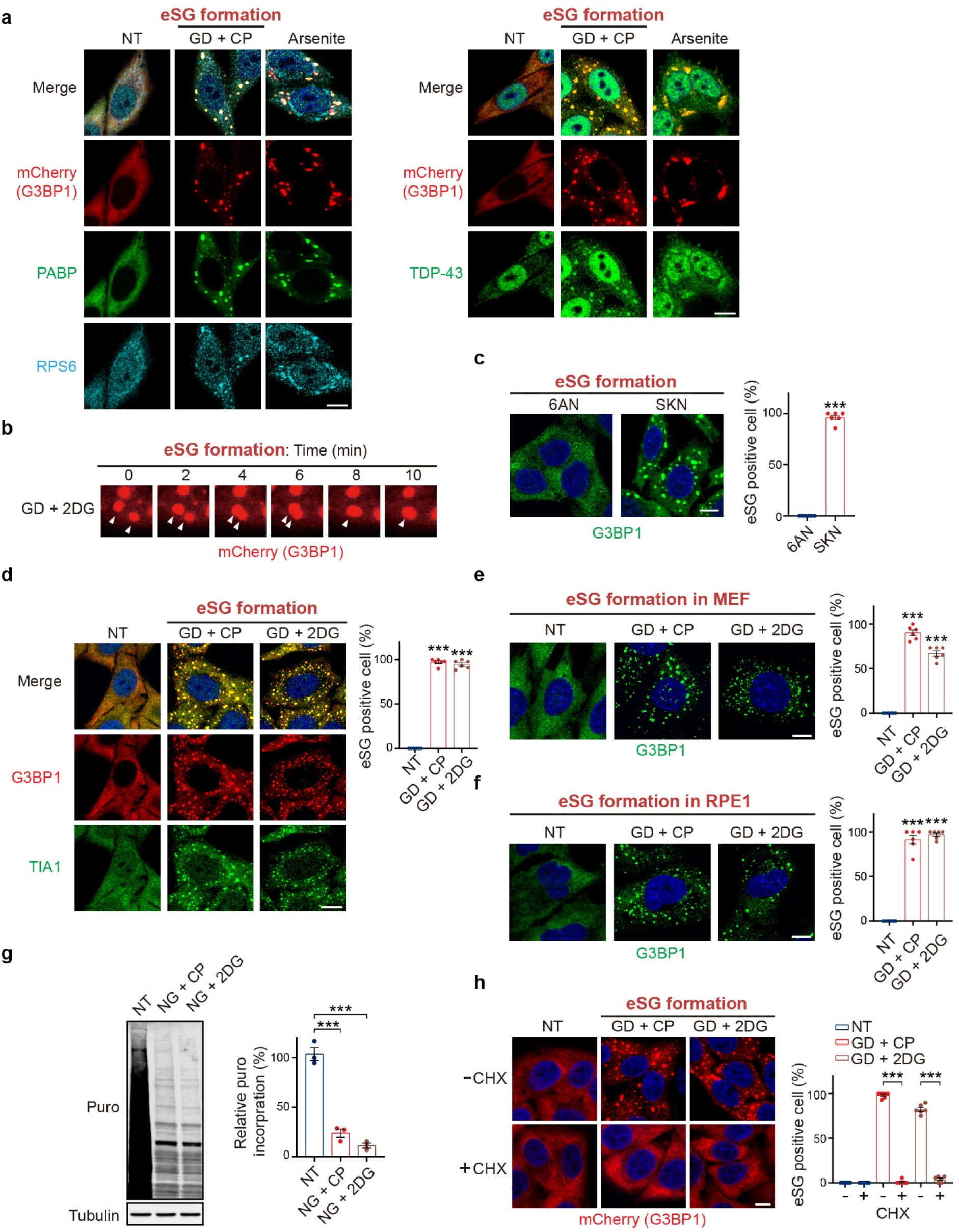
Inhibition of glycolysis, but not PPP, triggers eSG formation and translational inhibition and is dependent on polysome disassembly. **a**, Representative images of SGs or eSGs in HeLa cells stably expressing mCherry-G3BP1 with the indicated treatments. Cells were immunostained with anti-PABP, RPS6, and TDP-43 antibodies. **b**, A representative fusion of eSGs during glycolytic blockage-induced energy stress condition in an mCherry-G3BP1-expressing cell is shown. **c**,**d**, Representative images and quantification of eSGs formed in HeLa cells treated with 6AN (250 μM) or SKN (25 μM) (**c**) or the indicated energy stresses (**d**) for 1 h. The eSGs were revealed by immunofluorescence (IF) of G3BP1 (**c**) or G3BP1 combined with TIA1 (**d**) and assessed as the percentages of eSG positive cells (n = 6). **e**,**f**, Images of eSGs formed in MEF (**e**) or human RPE1 cells (**f**) after the indicated treatments for 1 h. eSGs were visualized and quantified by G3BP1 IF (n = 6). **g**, HeLa cells after 1 h of the indicated treatments were subjected to the ribopuromycinylation assay and immunoblotting to quantify the translational rates (n = 3). **h**, HeLa cells stably expressing mCherry-G3BP1 were treated with indicated energy stress for 1 h in the presence or absence of cycloheximide (CHX). eSGs were quantified as the percentages of eSG-positive cells (n = 6). Nuclei were visualized by DAPI staining (blue). Data are shown as means ± SEM, analyzed by unpaired two-sided Student’s *t*-test. * P < 0.05; ** P < 0.01; *** P < 0.001; ns, not significant. Scale bars, 10 μm.

**Extended Data Fig. 2.**
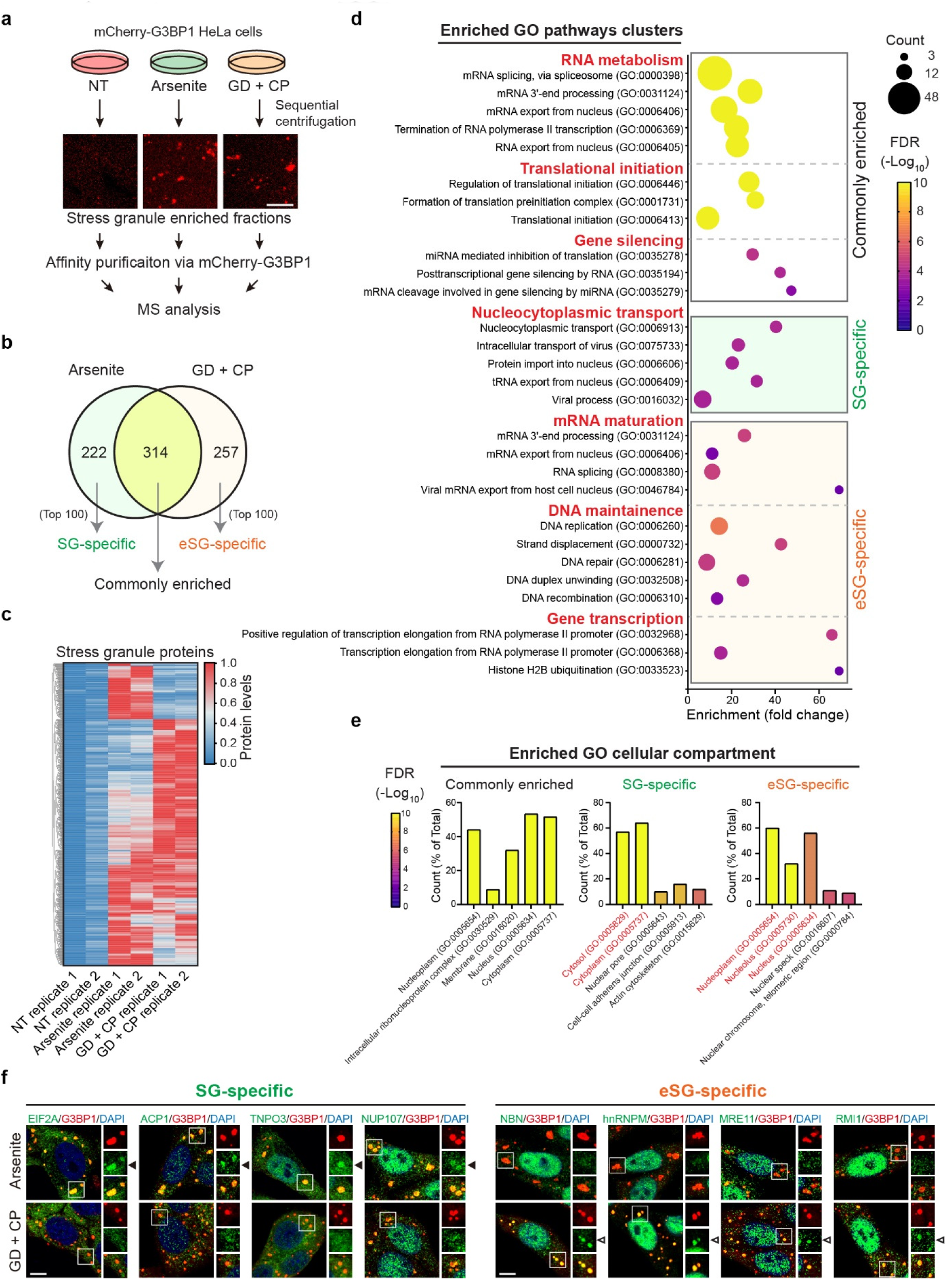
Protein compositions of SGs and eSGs. **a**, Scheme for the preparation of stress granule enriched fractions from HeLa cells stably expressing mCherry-G3BP1. **b**, Venn diagram depicting the number of the 793 granule proteins identified in either the SG or eSG fraction. **c**, Heat map showing the relative protein levels of the 793 granule proteins in either granule fraction as compared to the nontreated control. There were two biological replicates for each condition. **d**, GO biological pathway analysis of the SG- or eSG-enriched proteins and the proteins common to both. The GO terms of the significantly enriched pathways (FDR < 0.01), protein numbers, and fold changes in enrichment are given. The biological processes indicated by the clustered GO terms are listed and highlighted in red. **e**, GO cellular compartment analysis of the SG- or eSG-enriched proteins and those common to both. For each group, the top five GO terms (FDR) and percentage of proteins in each group are presented. The cytosolic GO terms of SG-specific proteins or nuclear GO terms of eSG-specific proteins are highlighted in red. **f**, HeLa cells were stressed with arsenite by glycolysis inhibition (GD+ CP) for 1 h to trigger SG or eSG, respectively. Stress granules were visualized by G3BP1 IF, and the granule localization of the four top-ranked SG-specific proteins (EIF2A, ACP1, TNPO3, or NUP107) or eSG-specific proteins (NBN, hnRNPM, MRE11, or RMI1) is shown. Closed arrowheads, proteins specifically localized to SGs; open arrowheads, proteins specifically localized to eSGs. Nuclei were visualized by DAPI staining (blue). Scale bars, 10 μm.

**Extended Data Fig. 3.**
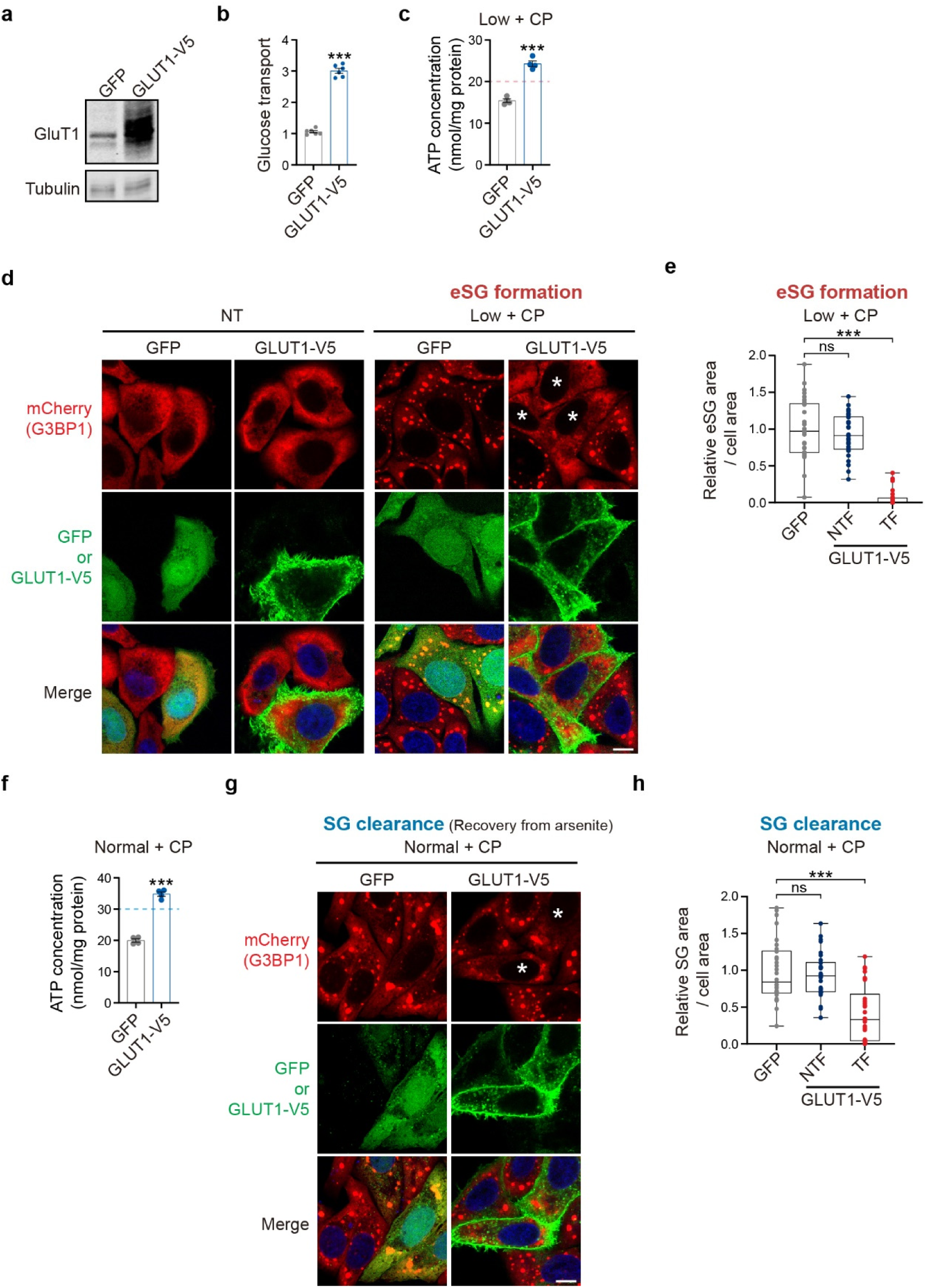
Overexpression of GLUT1 Diminishes eSG formation and restores the SG clearance in cells treated with glycogenolysis inhibition in limited glucose. **a**, Western blot analysis of GLUT1 levels in HeLa cells transiently overexpressing either GFP or V5-tagged GLUT1 (GLUT1-V5). **b**, Glucose transport assay of HeLa cells overexpressing GFP or GLUT1-V5 (n = 6). **c**, Intracellular ATP concentrations in HeLa cells overexpressing GFP or GLUT1-V5 and treated with severe energy stress induced by the inhibition of glycogenolysis in low physiological glucose (Low + CP) (n = 3). The ATP level below which eSG formation was triggered is indicated by dotted lines. **d**, Representative images of the eSGs triggered by the indicated treatments for 1 h in HeLa cells stably expressing mCherry-G3BP1 and transfected with either GFP or GLUT1-V5. Asterisks indicate cells with over-expressed GLUT1-V5 and diminished eSGs. **e**, Quantification of (**d**) as the relative eSG area per cell in cells transfected with either GFP (GFP) or GLUT1-V5 (TF) or in non-transfected (NTF) cells (n = 30 cells from three independent experiments). **f**, Intracellular ATP concentrations in HeLa cells overexpressing GFP or GLUT1-V5 and treated with moderate energy stress introduced by the inhibition of glycogenolysis in normal physiological glucose (Normal + CP) (n = 3). The ATP level below which SG clearance was impaired is indicated by dotted lines. **g**, Representative images of the persistent SGs after recovery from arsenite treatment for 1 h in the presence of the indicated energy stress in HeLa cells stably expressing mCherry-G3BP1 and transfected with either GFP or GLUT1-V5. Asterisks indicate cells with over-expressed GLUT1-V5 and a reduction in persistent SGs. **h**, Quantification of (**g**) as the relative area of persistent SGs per cell in GFP, NTF, or TF GLUT1-V5 cells (n = 30 cells from three independent experiments). Nuclei were visualized by DAPI staining (blue). Data are shown as means ± SEM, analyzed by unpaired two-sided Student’s *t*-test. * P < 0.05; ** P < 0.01; *** P < 0.001; ns, not significant. Scale bars, 10 μm.

**Extended Data Fig. 4.**
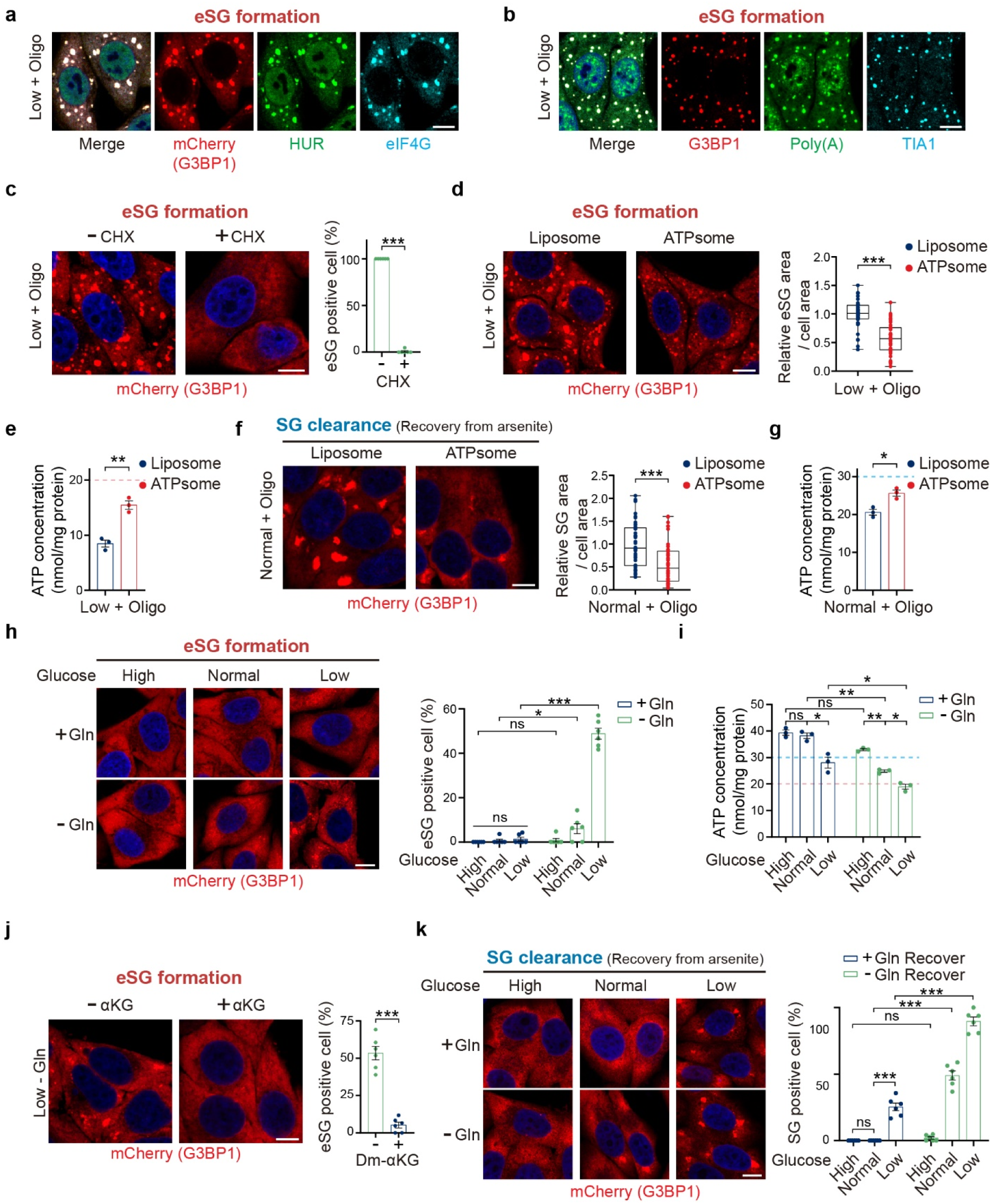
ATP reductions account for the OXPHOS-mediated eSG and SG dynamics. **a**,**b**, Representative images of eSGs in HeLa cells, with (**a**) or without (**b**) stable expression of mCherry-G3BP1 and treated by oligomycin administration in physiologically low glucose (Low + Oligo). Cells were stained with anti-HUR and anti-eIF4G antibodies (**a**) or anti-TIA1 antibody and mRNA FISH using an oligo-dT_20_ probe (**b**). **c**, HeLa cells stably expressing mCherry-G3BP1 were treated with the indicated energy stress in the presence or absence of cycloheximide. The eSGs were quantified as the percentage of eSG-positive cells (n = 6). **d**, Representative images and quantification of eSG formation, as assessed by the relative eSG areas per cell, in mCherry-G3BP1-expressing cells receiving the indicated treatment in the presence of either control empty liposomes or ATP-containing liposomes (ATPsome) (n = 45 cells from three independent experiments). **e**, Intracellular ATP concentrations in cells treated as in (**d**) (n = 3). The ATP level below which eSG formation was triggered is indicated by dotted lines. **f**, Representative images and quantification of persistent SGs in mCherry-G3BP1-expressing cells after recovery from arsenite treatment in physiologically normal glucose in the presence of OXPHOS inhibition (Normal + Oligo) plus either control liposomes or ATP-liposomes (ATPsome). SGs were quantified as the relative eSG areas per cell (n = 45 cells from three independent experiments). **g**, Intracellular ATP concentrations in cells treated as in (**f**) (n = 3). The ATP level below which SG clearance was impaired is indicated by dotted lines. **h**, Images of eSGs in G3BP1-expressing HeLa cells treated with the physiological range of glucose for 1 h in the presence (+Gln) or absence (-Gln) of glutamine. The eSGs were quantified as the percentage of eSG-positive cells (n = 6). **i**, Intracellular ATP concentrations in cells treated as in (**h**) (n = 3). The ATP levels below which eSG formation was triggered (pink) or SG clearance was impaired (cyan) are indicated by dotted lines. **j**, eSGs formation capacity in G3BP1-expressing HeLa cells triggered by glutamine deprivation in low glucose (Low - Gln) in the presence (+αKG) or absence (-αKG) of Dm-αKG. The eSGs were quantified as the percentage of eSG-positive cells (n = 6). **k**, Persistent SGs in mCherry-G3BP1 expressing cells after recovery for 1 h after arsenite removal in medium containing the physiological range of glucose, with or without glutamine deprivation. The SGs were quantified as the percentage of SG positive cells (n = 6). Nuclei were visualized by DAPI staining (blue). Data are shown as means ± SEM, analyzed by unpaired two-sided Student’s *t*-test. * P < 0.05; ** P < 0.01; *** P < 0.001; ns, not significant. Scale bars, 10 μm.

**Extended Data Fig. 5.**
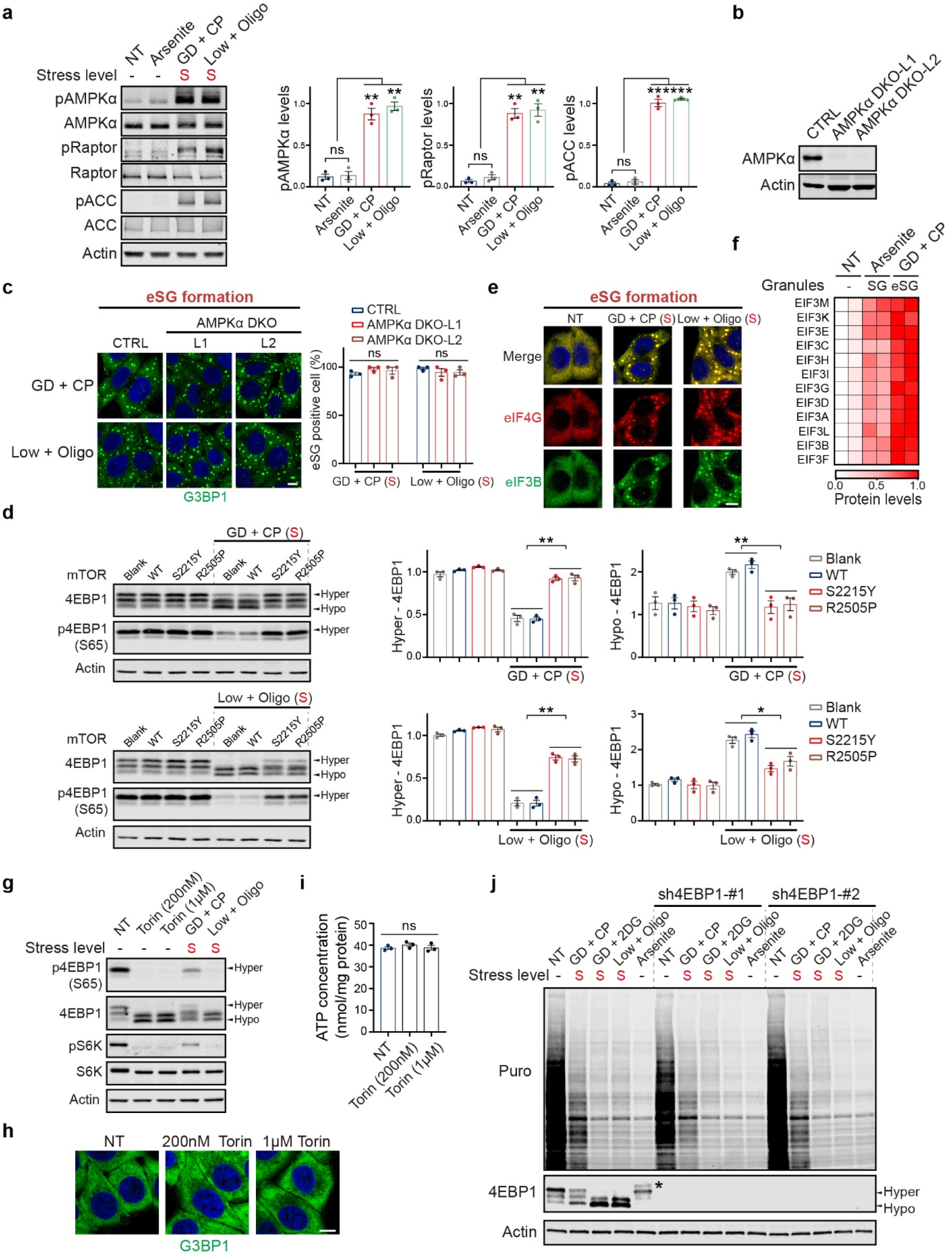
AMPK is not involved in eSG formation, and forced dephosphorylation of 4EBP1 is not sufficient to trigger eSG formation. **a**, Western blot analysis of AMPK activity using the indicated antibodies in cells treated with severe energy stress or arsenite. The relative abundance of phosphorylated AMPK (pAMPKα, Thr172), Raptor (pRaptor, Ser792), or ACC (pACC Ser79) was normalized to the maximal level of each target (n = 3). **b**, Western blotting confirms the deletion of AMPKα in two independent HeLa cell lines lacking both the α1 and α2 subunits of AMPK (AMPKα DKO-L1, L2). CTRL: control cells expressing a blank vector. **c**, Representative images of eSGs in the control or the two AMPK DKO lines with the indicated treatments. eSGs were quantified as the percentage of eSG-positive cells (n = 3). **d**, Cells overexpressing wild-type (WT) or mutant mTOR proteins or a blank vector (CTRL) were subjected to severe energy stress treatments and western blotting analysis using the indicated antibodies. Hypophosphorylated and hyperphosphorylated 4EBP1 (hypo- and hyper-4EBP1) are indicated. The hyper-4EBP1 levels were assessed by Ser65 phosphorylation, and the hypo-4EBP1 levels were assessed by the relative proportion of the lowest band of 4EBP1 to the total protein (n = 3). **e**, Localization of eIF3b in the eSGs marked by eIF4G in cells receiving the indicated treatments. **f**, Heat map showing the relative protein levels of 12 eIF3 proteins identified in the granule fractions selected from (Extended Data Fig. 2c). **g**, Western blot analysis of the hyper- and hypo-4EBP1 and pS6K levels in HeLa cells treated with Torin1 at the indicated concentrations or with severe energy stress for 1 h. **h**,**i**, Images of G3BP1 IF and the intracellular ATP concentrations (n = 3) in HeLa cells treated with Torin1 as in (**g**). **j**, HeLa cells with or without 4EBP1 depletion were treated with arsenite or the indicated severe energy stress for 1 h and then subjected to the ribopuromycinylation assay and immunoblotting to reveal the translational rates. The asterisk indicates a band of 4EBP1 that appears after arsenite treatment as a result of unidentified modifications. Nuclei were visualized by DAPI staining (blue). The stress levels of each treatment are indicated. S, severe energy stress. Data are means ± SEM, analyzed by unpaired two-sided Student’s *t*-test. * P < 0.05; ** P < 0.01; *** P < 0.001; ns, not significant. Scale bars, 10 μm.

**Extended Data Fig. 6.**
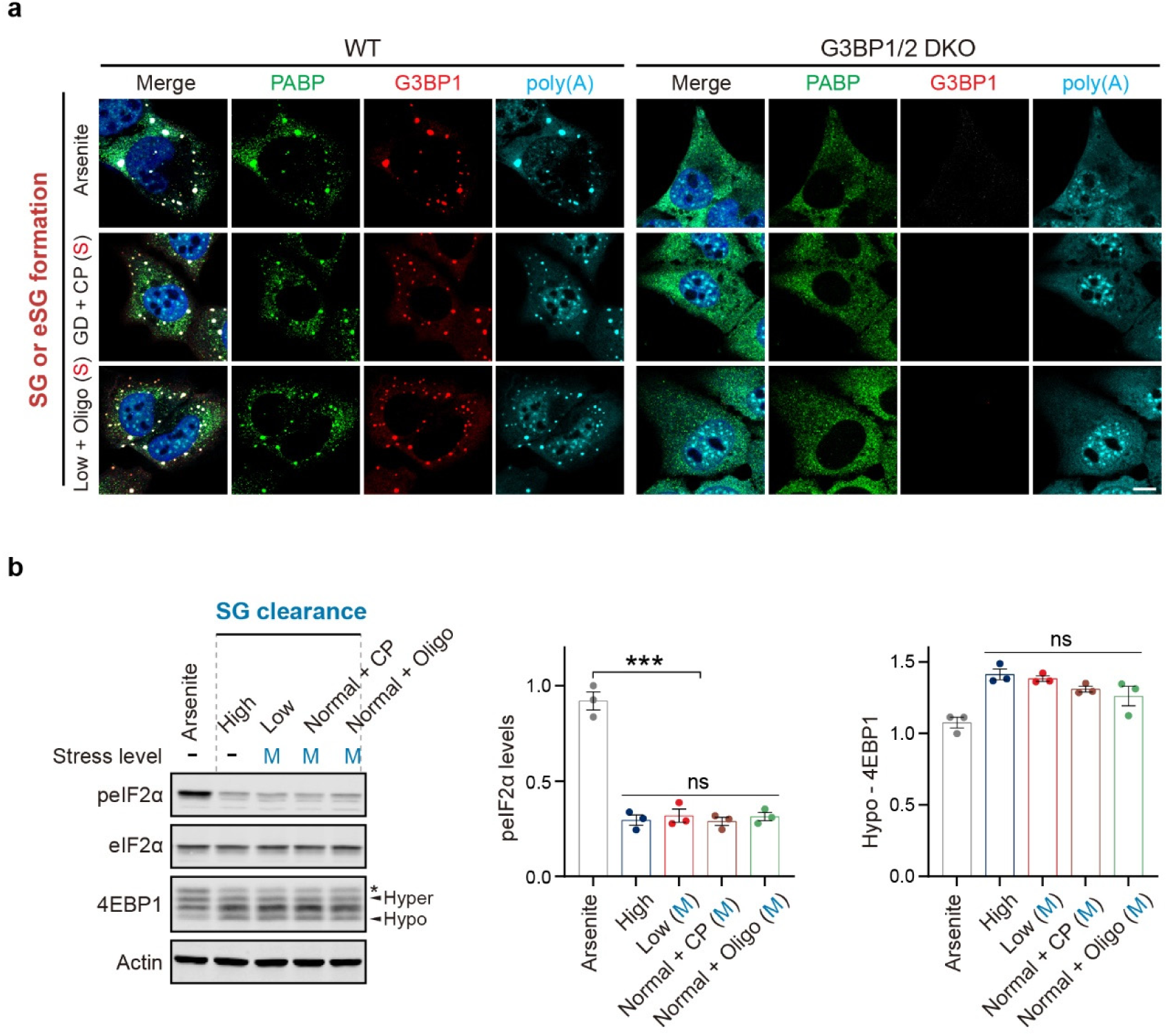
The absence of eSG formation in cells lacking G3BP and unchanged peIF2a and p4EBP1 levels under moderate energy stress conditions. **a**, Representative images of SGs or eSGs in wild-type or G3BP1/G3BP2 double-knockout (DKO) human U2OS cells. Cells receiving the indicated treatments for 1 h were immunostained with anti-PABP and anti-G3BP1 antibodies, and subjected to mRNA FISH using an oligo-dT20 probe. **b**, Western blotting analysis and quantifications of the peIF2a and hypo-4EBP1 levels in cells after the indicated treatments. Hyper- and hypo-4EBP1 are indicated. The asterisk indicates a band of 4EBP1 that appears after arsenite treatment because of some unidentified modification. There was no difference in the eIF2a or 4EBP1 pattern between the cells in high glucose and the cells with moderate energy stress. The stress levels of each treatment are indicated. M, moderate energy stress. Data are shown as means ± SEM, analyzed by one-way ANOVA. *** P < 0.001; ns, not significant. Scale bars, 10 μm.

**Extended Data Fig. 7.**
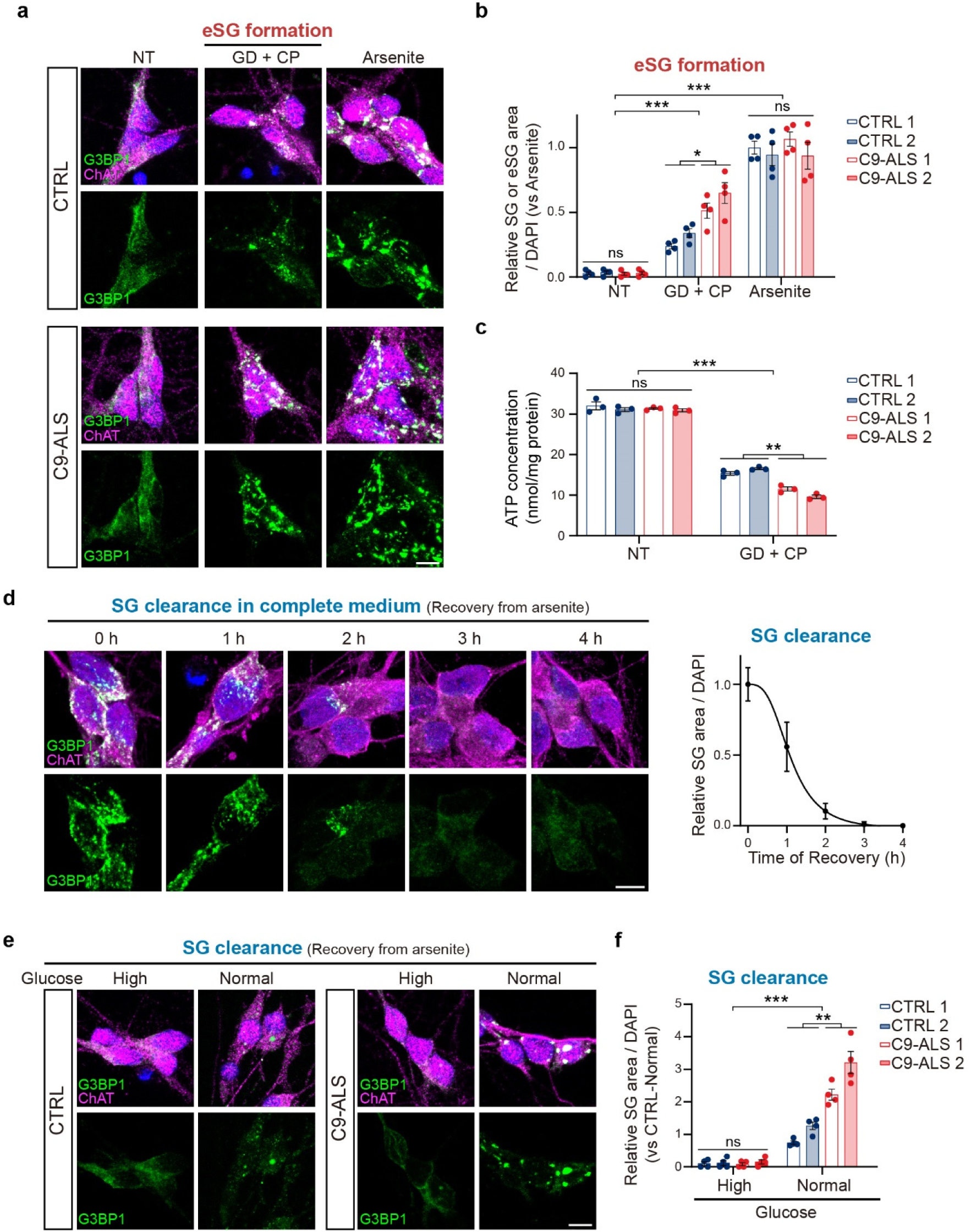
Disrupted eSG formation and SG clearance in C9-ALS patients’ motor neurons after energy stress. **a**,**b**, Two healthy control and two C9-ALS motor neurons (MNs) treated with either arsenite or the indicated severe energy stress for 1 h were subjected to immunostaining analysis with G3BP1 and a motor neuron marker, ChAT. Representative images of MNs (CTRL 1 and C9-ALS 1) are shown (**a**). The quantification of SGs or eSGs in MNs was assessed by the relative granule area per cell (**b**) (n = 4). **c**, Intracellular ATP concentrations in healthy control and C9-ALS MNs with or without the severe energy stress treatment as in (**a**) (n = 3). **d**, Healthy human MNs treated with arsenite for 1 h to trigger SG formation were allowed to recover in complete medium for the indicated times. The persistent SGs in MNs were visualized by G3BP1 and ChAT IF and quantified as the relative SG areas per cell and normalized to the value at time point 0 (n = 3). Data are plotted with the time points. **e**,**f**, Healthy control and C9-ALS MNs were allowed to recover from arsenite treatment in mediums with the indicated glucose concentration for 3 h. MNs were subjected to immunostaining analysis with G3BP1 and ChAT IF. Representative images of MNs (CTRL1 and C9-ALS1) are shown (**e**). The quantification of persistent SGs in two healthy control or two C9-ALS MNs in physiologically high or normal glucose is shown in (**f**) (n = 4). Nuclei were visualized by DAPI staining (blue). Data are means ± SEM, analyzed by unpaired two-sided Student’s *t*-test. * P < 0.05; ** P < 0.01; *** P < 0.001; ns, not significant. Scale bars, 5 μm.

**Supplementary Table 1. Proteomic analyses of SG and eSG components.**

(see the supplementary document)

## Notes

### Competing Interest Statement

The authors have declared no competing interest.

